# Decisions bias future choices by modifying hippocampal associative memories

**DOI:** 10.1101/802462

**Authors:** Lennart Luettgau, Claus Tempelmann, Luca Franziska Kaiser, Gerhard Jocham

## Abstract

Decision making is guided by memories of option values. However, retrieving items from memory renders them malleable. Here, we show that merely retrieving values from memory and making a choice between options is sufficient both to induce changes to stimulus-reward associations in the hippocampus and to bias future decision making. After allowing participants to make repeated choices between reward-conditioned stimuli, in the absence of any outcome, we observed that participants preferred stimuli they had previously chosen, and neglected previously unchosen stimuli, over otherwise identical-valued options. Using functional brain imaging, we show that decisions induced changes to hippocampal representations of stimulus-outcome associations. These changes were correlated with future decision biases. Our results indicate that choice-induced preference changes are partially driven by choice-induced modification of memory representations and suggest that merely making a choice - even without experiencing any outcomes - induces associative plasticity.

## Introduction

According to neo-classical economic models of decision making, choices are guided by memories of option values (Ariely & Norton, 2008). However, this unidirectional view has been challenged by cognitive accounts of decision making, suggesting that memory representations of option values might themselves be subject to changes induced by an agent’s choices. This suggests a bidirectional relationship between value representations in memory and decision making (Ariely & Norton, 2008; Riefer, Prior, Blair, Pavey, & Love, 2017).

Even though real-life decisions often involve memory retrieval of learned associations between reward-predictive cues and outcomes, memory mechanisms underlying choice-induced preference changes have, to the best of our knowledge, never been systematically studied (Ariely & Norton, 2008; Brehm, 1956; Schonberg et al., 2014; Sharot, Velasquez, & Dolan, 2010). This might be partially related to the fact that most studies on choice-induced preference changes employed direct presentation of the outcomes to be chosen. However, this approach by design obliterates and confounds underlying associative learning contributions to the revaluation process (Izuma et al., 2010), and is blind to related memory processes, such as retrieval competition (Wimber, Alink, Charest, Kriegeskorte, & Anderson, 2015).

In naturalistic decision making scenarios, choices often have to be made without direct experience of feedback or rewards. Instead, decision makers have to rely on relational knowledge of stimuli, actions and outcomes. The absence of direct external feedback and the resulting inability to adjust synaptic weights based on error-driven learning mechanisms suggests the use of unsupervised learning to optimize behavior in those situations. Likely candidate mechanisms for such unsupervised behavioral adaptation are memory retrieval dynamics. It is well established that retrieval of an item from memory, e.g. a conditioned stimulus (CS) triggering retrieval of an associated outcome, leads to improved remembering. However, memory for competing items, e.g. a CS associated with the same outcome, is impaired simultaneously (Anderson, Bjork, & Bjork, 1994; Hulbert & Norman, 2015; Wimber et al., 2015). Such retrieval-induced forgetting (Anderson et al., 1994) would predict choice biases towards previously chosen CS based on retrieval-related strengthening of CS-US association. However, retrieval-induced forgetting would predict the same effect for a previously presented, but unchosen CS, since both chosen and unchosen CS activate neural populations representing the respective associated outcome (Barron, Dolan, & Behrens, 2013; Boorman, Rajendran, O’Reilly, & Behrens, 2016; Howard, Kahnt, & Gottfried, 2016; Klein-Flugge, Barron, Brodersen, Dolan, & Behrens, 2013; Onat & Büchel, 2015; Tonegawa, Morrissey, & Kitamura, 2018). A more recent theoretical framework (Nonmonotonic Plasticity Hypothesis, as reviewed in Ritvo, Turk-Browne, & Norman, 2019) suggests U-shaped spreading activation during associative memory retrieval: Inactive memories remain unaltered, whereas moderately activated associative memories are weakened, and higher activation leads to strengthening of memories (Ritvo et al., 2019). Translating this idea to memory-based decisions between two CS, we assumed that both CS would moderately activate neural populations representing the associated outcome (as the outcome is never presented). However, consistent with the finding that chosen options receive higher attentional weighting than unchosen options (as reflected in higher learning rates for chosen options (Klein, Ullsperger, & Jocham, 2017; Palminteri, Khamassi, Joffily, & Coricelli, 2015)), we further assumed that choices of a CS would induce additional activation of the associated outcome, whereas this would not be the case for unchosen CS, retaining an intermediate activation state of the associated outcome. Thus, we hypothesized that choosing a CS would strengthen the related stimulus-outcome association. Contrarily, not choosing a CS would weaken the respective stimulus-outcome association. We expected that these choice-induced alterations of the associative memory structure would result in subsequent preference changes. In other words, we assumed that choices themselves can act as self-generated “teaching signals”, dynamically altering stimulus-outcome associations stored in memory by shifting associative memories along a nonmonotonic plasticity function (Ritvo et al., 2019).

We expected choice-induced preference changes to be driven by modifications of stimulus-outcome associations in the hippocampus and lateral orbitofrontal cortex, two key regions for storing and updating associative representations (Boorman et al., 2016; Klein-Flugge et al., 2013).

Thus, the goal of the present study was twofold. First, we aimed at investigating how choice-related alterations of associative memories bias future decision making. Second, we sought to investigate a neurobiologically plausible mechanism underlying choice-induced preference changes. To test our key predictions, we designed a novel learning and decision making paradigm which we used in three independent behavioral experiments and one functional magnetic resonance imaging (fMRI) experiment. For the fMRI study, we exploited repetition suppression (RS) effects (Barron et al., 2013; Barron, Garvert, & Behrens, 2016; Garvert, Dolan, & Behrens, 2017; Grill-Spector & Malach, 2001; Klein-Flugge et al., 2013) to measure associative strength between conditioned (CS) and unconditioned stimuli (US) (Boorman et al., 2016; Klein-Flugge et al., 2013). Participants first established Pavlovian associations between CS and differently valued US. Next, in a choice-induced revaluation, participants made binary choices between differently valued CS, without observing the associated US. Finally, in a probe phase, where they made choices between all possible CS combinations, participants showed preference increases for previously chosen, and preferences decreases for previously unchosen CS, compared to otherwise equivalent CS. These choice-induced alterations in decision behavior were accompanied by corresponding changes in CS-US RS effects in the hippocampus and lateral orbitofrontal cortex. These findings were corroborated by multivariate pattern similarity analyses (a variant of representational similarity analysis, RSA (Kriegeskorte, Mur, & Bandettini, 2008)). Furthermore, the magnitude of the hippocampal RS effect was correlated with individual probe phase decision biases.

## Results

### Behavioral Experiments

First, we detailed the behavioral choice-induced revaluation effect in three independent experiments. In each experiment, participants learnt associations between neutrally rated CS and three food items (Blechert, Meule, Busch, & Ohla, 2014) serving as unconditioned stimuli (US, Fig. 1A). For each participant, the US were individually chosen based on a prior rating of subjective preference on a visual analog scale ranging from low to high preference. This rating procedure resulted in selection of a low-value (US^−^), an intermediate-value (US^0^) and a high-value (US^+^) food item. Next, participants rated kanji stimuli according to their subjective preference. Six of these kanjis rated in close proximity to “neutral” (center point of the visual analog scale) were selected as CS for Pavlovian learning. Two CS each were paired with one US, resulting in three categories of differently valued CS: CS^+^_A/B_, CS^0^_A/B_ and CS^−^_A/B_, for high-, intermediate- and low-value CS.

**Fig. 1.**
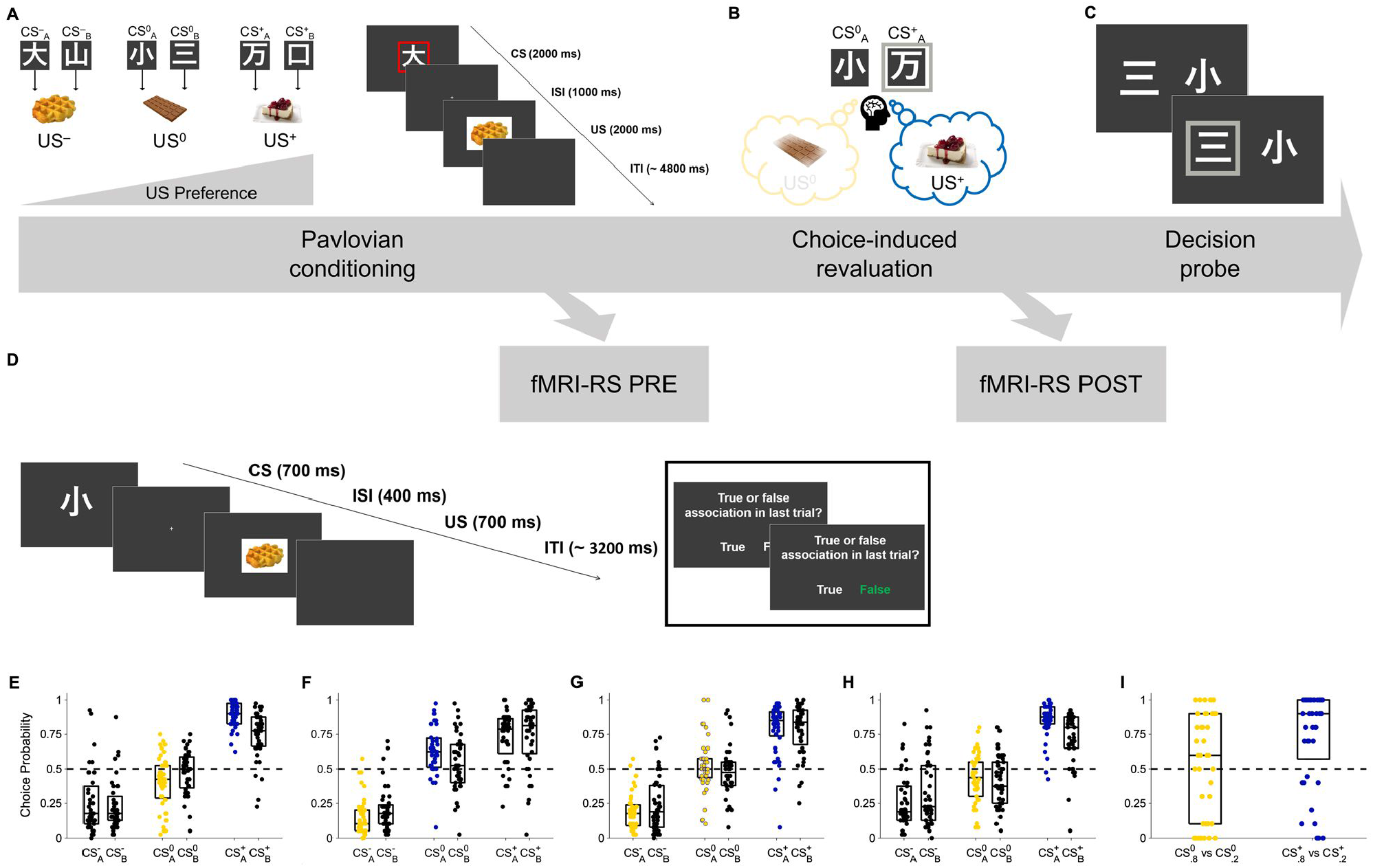
Task schematic and behavioral results. A) Participants rated subjective desirability of conditioned stimuli (CS, kanjis) and unconditioned stimuli (US, food items). During Pavlovian conditioning, participants learned to associate six CS with three US. Each US was associated with two CS. B) Choice-induced revaluation: After Pavlovian conditioning, participants made choices between CS^+^_A_ versus CS^0^_A_ (Experiments 1 & 4), CS^0^_A_ versus CS^-^_A_ (Experiment 2), CS^0^ _A_ versus either CS^-^_A_ or CS^+^_A_ (Experiment 3), and CS^+^_80_ versus CS^0^_80_ or CS^+^_20_ versus CS^0^_20_ (Experiment 5), interleaved with lure decisions. C) Decision probe: Following choice-induced revaluation, participants made binary choices between all possible combinations of CS, or CS^+^_80_ versus CS^+^_20_ and CS^0^_80_ versus CS^0^_20_ (Experiment 5), to assess preferences. D) Attentional control task performed during fMRI repetition suppression (Experiment 4). E-H) Previously chosen CS (blue scatter) are selected more often compared to equivalent CS (black scatter) in Experiments 1, 2, 4 (E, F, H) and previously unchosen CS (yellow scatter) are selected less often compared to equivalent CS (black scatter) in Experiment 1, 2 (E, F) during decision probe. The effect is not present in Experiment 3 (G), indicating that the roughly equal proportion of choices and non-choices of CS^0^_A_ during revaluation had cancelled each other out. I) Behavioral control experiment (Experiment 5), orthogonalizing contributions of “go” and “no-go” tagging and associative strength between CS and US to choice probabilities. Previously chosen (“go tag”) and strongly associated CS^+^_80_ is preferred over previously chosen and weakly associated CS^+^_20_ (blue scatter), while there is only descriptive evidence for preference of previously unchosen (“no-go tag”) and strongly associated CS^0^_80_ over previously unchosen and weakly associated CS^0^_20_.

The Pavlovian learning phase was followed by a choice-induced revaluation. From two value categories, one CS each was selected, and participants made binary choices between them, interspersed with lure decisions between non-reward-predictive kanjis (Fig. 1B). Crucially, no associated US were presented following choices, excluding the possibility of alterations in strength of stimulus-outcome associations due to directly experienced outcomes. The choice-induced revaluation phase was followed by a decision probe phase in which participants chose repeatedly between all binary CS combinations to assess preferences (Fig. 1C). The key comparison was between CS presented during revaluation versus CS from the same value category that had not been presented. Again, no outcomes were presented.

### Decisions are biased by past choices

There was evidence for value transfer from US to CS across all studies (Fig. 1E-H), as indicated by significant main effects of CS value on probe phase choice probabilities across all four experiments (all *F*s > 94.99, *P*s < .001, η^2^_*p*_s > .69, 1–*β*s > .99, repeated-measure analyses of variance, rmANOVA). Decision making during the revaluation phase had clearly dissociable effects on choices during the probe phase. CS that were chosen during the revaluation phase were more likely to be selected during the later probe phase compared to the CS of equal value that were not presented during choice-induced revaluation.

In Experiment 1, participants (*N* = 40) made choices between the intermediate-value CS^0^_A_ and the high-value CS^+^_A_ during the choice-induced revaluation phase. As we had directed hypotheses for the choice effects (increased choice probabilities for the previously chosen and decreased choice probabilities for the previously unchosen CS), we used one-tailed post hoc tests.

In the probe phase, participants preferred CS^+^_A_, the previously selected stimulus, compared to CS^+^_B_ (*Z* = 3.98, *P* < .001, Cohen’s *U*_*3*_ *=* .*85* (range 0 – 1, .5 indicating no effect), Wilcoxon signed-rank test, one-tailed). This effect was mainly driven by preference for CS^+^_A_ in pairwise within-category choice trials between CS^+^_A_ and CS^+^_B_ (*Z* = 3.43, *P* < .001, Cohen’s *U3*_*1*_ *=* .69 (range 0 – 1, .5 indicating no effect), one-sample Wilcoxon signed-rank test vs. 0.5, one-tailed; Supplementary Fig. S2E). Conversely, participants were less likely to select CS^0^_A_, the previously non-selected stimulus, compared to CS^0^_B_ (*Z* = 1.97, *P* = .025, *U*_*3*_ *=* .70, Wilcoxon signed-rank test, one-tailed). Again, this effect was mainly driven by reduced choice of CS^0^_A_ in pairwise within-category choice trials directly contrasting CS^0^_A_ and CS^0^_B_ (*Z* = 2.05, *P* = .020, *U3*_*1*_ = .68, one-sample Wilcoxon signed-rank test vs. 0.5, one-tailed; Supplementary Fig. S2E). The observed dissociation in choice behavior was also evident in a significant interaction effect of CS value × CS type (A or B): *F*_*2, 78*_ = 10.01, *P* < .001, η^2^_*p*_ = .20, 1–*β* > .99 (rmANOVA, Fig. 1E). Thus, compared to otherwise equivalent CS, participants exhibited a systematic preference for CS they had previously chosen, whereas they displayed a diminished preference of CS they had previously not chosen.

After having established that an intermediate value CS (CS^0^_A_) could be devalued by non-choices, we next asked in Experiment 2, whether we can induce the exact opposite, an increase in preference for CS^0^_A_. Therefore, participants (*N* = 40) were presented with decisions between intermediate-value CS^0^_A_ and low-value CS^−^_A_ during the choice-induced revaluation phase. Conceptually replicating the results of Experiment 1, participants in Experiment 2 favored the previously chosen CS^0^_A_ over CS^0^_B_ (*Z* = 2.20, *P* = .014, *U*_*3*_ = .68, Wilcoxon signed-rank test, one-tailed) during the decision probe phase, resulting from preference for CS^0^_A_ in pairwise within-category choice trials between CS^0^_A_ and CS^0^_B_ (*Z* = 1.93, *P* = .027, *U3*_*1*_ *=* .68, one-sample Wilcoxon signed-rank test vs. 0.5, one-tailed; Supplementary Fig. S2F). Contrarily, participants neglected CS^−^_A_ overall compared to CS^−^_B_ (*Z* = 1.91, *P* = .028, *U*_*3*_ *=* .*66*, Wilcoxon signed-rank test, one-tailed), resulting from descriptively reduced preference for CS^−^_A_ in pairwise within-category choice trials between CS^−^_A_ and CS^−^_B_ (*Z* = 1.41, *P* = .079, *U3*_*1*_ = .63, one-sample Wilcoxon signed-rank test vs. 0.5, one-tailed; Supplementary Fig. S2F). Again, there was a significant interaction effect of CS value × CS type (*F*_*2, 78*_ = 4.84, *P* = .01, η^2^_*p*_ = .11, 1–*β* > .99, rmANOVA, Fig. 1F) indicating clearly dissociable choice behavior during the decision probe. This pattern of results suggests a ubiquitous, value-independent mechanism of choice-induced revaluation.

These results so far show that choices and non-choices act in opposite directions. Consequently, we predicted that it would be possible to cancel out choice-induced preference increases and devaluation of CS^0^_A_. To test this prediction, in Experiment 3 (*N* = 44), we presented an equal amount of binary decisions between CS^0^_A_ and CS^-^_A_ as between CS^0^_A_ and CS^+^_A_ during choice-induced revaluation. Since choice-induced revaluation effects for CS^0^_A_ should cancel each other out, unlike in Experiment 1 and 2, we did not have directed hypotheses for the choice effects. We thus used two-tailed tests for the corresponding post hoc tests. As expected, there was no evidence for change in preference for CS^0^_A_ compared to CS^0^_B_ (*Z* = 0.41, *P* = .68, *U*_*3*_ = .55, Wilcoxon signed-rank test, two-tailed, Fig. 1G). Consistently, there was no evidence for preference changes for CS^0^_A_ in pairwise within-category choice trials between CS^0^_A_ and CS^0^_B_ (*Z* = 0.12, *P* = .905, *U3*_*1*_ = .57, one-sample Wilcoxon signed-rank test vs. 0.5, two-tailed; Supplementary Fig. S2G), suggesting that the effects of choices and non-choices had indeed cancelled each other out (interaction effect of CS value × CS type: *F*_2, 86_ = 1.31, *P* = .280, η^2^_*p*_ = .03, 1–*β* = .71, rmANOVA).

Importantly, the dissociations observed in choice behavior in Experiment 1 & 2 rule out alternative accounts for explaining choice behavior, such as extinction or mere exposure effects. Both accounts would predict unidirectional preference changes for the CS presented during choice-induced revaluation, independent of the choices made (decreases or increases in preference, respectively), which is incompatible with the present results.

However, an alternative explanation for the observed choice pattern is that participants learned simple choice rules for the two CS presented during the revaluation phase, akin to “go tags” for chosen CS (“choose this stimulus”) and “no-go tags” for unchosen CS (“do not choose this stimulus”). Accordingly, the observed changes in preferences could be attributed to simply repeating such choice heuristics acquired during revaluation. An additional behavioral experiment (Experiment 5, *N* = 40) was specifically designed to address this possibility. We orthogonalized contributions of associative strength and choice rule by letting participants assign choice-induced “go tags” to two chosen CS^+^ that differed in their associative strength to US^+^ (80% vs. 20% association) and “no-go tags” to two unchosen CS^0^ that likewise differed in their associative strength to US^0^ (80% vs. 20%). According to our hypothesis (“associative account”), probe phase decisions are guided by the learned associations and the strengthening/weakening of this association during revaluation. Therefore, we expected (after revaluation) a significantly increased choice probability for both highly associated stimuli: CS^0^_80_ should be preferred over CS^0^_20_, and CS^+^_80_ should be preferred over CS^+^_20_. On the contrary, if choice behavior was exclusively driven by learned “go” and “no-go tags” (“heuristic account”), and participants had learned “go tags” for both chosen CS^+^_80_ and CS^+^_20_ and “no-go tags” for both unchosen CS^0^_80_ and CS^0^_20_ (instead of changing the underlying associative structure), both same-value pairwise choice probabilities should be at chance level (CP = 0.50). Due to the directionality of our hypothesis, we used one-tailed tests.

Importantly, there was no significant difference between revaluation choice probabilities of CS^+^_80_ versus CS^0^_80_ and CS^+^_20_ versus CS^0^_20_ (*Z* = 1.19, *P* = .234, *U*_*3*_ = .53, Wilcoxon signed-rank test, two-tailed), ruling out unequal assignment of “go” and “no-go” tags across CS pairs. We observed that participants strongly favored the previously chosen and strongly associated CS^+^_80_ over the previously chosen and weakly associated CS^+^_20_ (*Z* = 3.55, *P* < .001, *U3*_*1*_ = .75, 1–*β* > .99, one-sample Wilcoxon signed-rank test, one-tailed). This pattern of results favors an explanation based on associative strengthening of the memory trace between CS^+^ and US^+^, rather than merely on expressing a “go tag”. Descriptively, participants also tended to favor the previously unchosen and strongly associated CS^0^_80_ compared to the previously unchosen and weakly associated CS^0^_20_ (*Z* = 0.61, *P* = .271, *U3*_*1*_ = .61, 1–*β* = .23, one-sample Wilcoxon signed-rank test, one-tailed, Fig. 1I) during the decision probe phase. This pattern of results favors an explanation based on associative strengthening of the memory trace between CS^+^ and US^+^, rather than merely on applying a choice heuristic. However, the non-significant preference for CS^0^_80_ over CS^0^_20_ does not provide definitive evidence against the alternative explanation that participants learned a “No-Go” tag for the unchosen stimulus. The observed asymmetry in the expression of response tendencies might result from differential mechanisms driving acquisition of “go” and “no-go” choice rules, akin to well-described Pavlovian biases(Guitart-Masip et al., 2012). Presumably, in high-value (CS^+^_80_ vs CS^+^_20_) choice trials, the majority of participants used the learned and updated CS-US associative strength instead of “go” response tendencies to guide their decisions, while this only tended to be the case for intermediate-value (CS^0^_80_ vs CS^0^_20_) choices (*Median* choice probability = .60, *range* = 0 – 1). Consistent with modelling and empirical evidence for asymmetric instrumental learning of action and inaction(Swart et al., 2017), we assume that reverting the initially learned action tendency for CS^+^_20_ could have less of an impeding effect on re-acquisition of a “no-go” response during decision probe phase, than re-acquisition of a “go” response for CS^0^_80_, which was initially learned with an inaction choice rule.

We fit six different variants of a reinforcement learning model using Rescorla-Wagner-like delta update rules (Rescorla & Wagner, 1972) for the four main experiments (Experiment 1-4). For each experiment, we compared the six models that implemented different ways by which participants could have learned CS-US associative strength - and updated associative strength during choice-induced revaluation based on “fictive” reward prediction errors. The “fictive” reward prediction errors were based on our reasoning that presentation of CS during revaluation would lead to retrieval of the associated US and that, consistent with our hypothesis, stimulus-outcome association of the chosen CS would be strengthened, whereas the association of the unchosen CS would be weakened. As no objective feedback (US) was presented during choice-induced revaluation, we assumed that the “fictive” reward prediction error would be computed as the difference between the CS value during previous presentation of the respective CS and subjective value of the current trial’s associated US, retrieved from associative memory. Our behavioral results were best captured by a computational model that differentially updated the learned CS-US associative strengths using “fictive” reward prediction errors elicited by revaluation phase decisions (Methods and Supplementary Table S1). Computational models using the best-fitting parameters successfully reproduced the observed empirical choice pattern (with the exception of the observed reduced choice probability of CS^0^_B_ in Experiment 4, Supplementary Figure S1 E-H). Thus, the best fitting models likely incorporate candidate computational mechanisms underlying the observed choice biases.

### Choices modify univariate neural measures of stimulus-outcome associations

Having established and replicated the behavioral effect of choice-induced revaluation in three independent behavioral samples, we next tested whether decisions induce changes of neural representations of CS-US associations.

In Experiment 4, we used functional magnetic resonance imaging and leveraged repetition suppression (RS) effects (Barron et al., 2013, 2016; Garvert et al., 2017; Grill-Spector & Malach, 2001; Klein-Flugge et al., 2013) to measure CS-US associative strength (Boorman et al., 2016; Klein-Flugge et al., 2013). When a neural ensemble is activated twice in brief succession (e.g. by rapid sequential presentation of the same visual stimulus), the second stimulus causes a diminished response. Accordingly, after learning the association between CS and US, the CS should elicit a representation of its associated US. Thus, presentation of the US itself, following the CS, should induce a diminished neural response. If the association between CS and US has been weakened by non-choices during revaluation, the CS is no longer capable of evoking the US representation to the same degree and should therefore elicit a stronger response (less repetition suppression). The same logic in reverse applies when the association has been strengthened by choices during revaluation.

As in the three behavioral experiments, participants first learned the six CS-US associations during Pavlovian learning. Following Pavlovian learning, we administered two RS blocks, one immediately before (“PRE”) and one immediately after (“POST”) the choice-induced revaluation phase, where participants (*N* = 42) made binary choices between CS^0^_A_ and CS^+^_A_. For the repetition suppression effects we had directed hypotheses (increased repetition suppression for the previously chosen, and decreased repetition suppression for the previously unchosen CS). Therefore, we used one-tailed post hoc tests. For the PRE phase, as well as for the control RS effects of CS^−^_A_ relative CS^−^_B_, there was no such directed hypothesis and we used two-tailed tests accordingly. Consistent with our hypothesis, we observed both a decrease in repetition suppression for CS^0^_A_ -US^0^ relative to CS^0^_B_ -US^0^, and an increase in repetition suppression between CS^+^_A_ -US^+^ compared to CS^+^_B_ -US^+^ (Fig. 2A, D) during POST but not during PRE (Fig. 2D inset, *Z* = 2.53, *P* = .006, *U*_*3*_ = .67, Wilcoxon signed-rank test, one-tailed) in the left hippocampus (Fig. 2A). Detailed analyses showed that this conjunction effect was driven by a dissociable effect (interaction effect CS value and time (PRE or POST), *F*_1, 41_ = 4.51, *P* = .040, η^2^_*p*_ = .10, 1–*β* = .99, rmANOVA): we found decreased repetition suppression for CS^0^_A_ (*Z* = 2.26, *P* = .012, *U3*_*1*_ = .67, one-sample Wilcoxon signed-rank test, one-tailed) and increased repetition suppression for CS^+^_A_ (*Z* = 2.26, *P* = .012, *U3*_*1*_ = .69, one-sample Wilcoxon signed-rank test, one-tailed) during POST, without any evidence for non-zero differences in PRE (all *Zs* < 1.05, *Ps* > .296, *U3*_*1*_ < .62, one-sample Wilcoxon signed-rank tests, two-tailed) or for the control contrast of either CS^−^_A_ or CS^−^_B_ (all *Zs* < 0.31, *Ps* > .760, *U3*_*1*_ < .55, one-sample Wilcoxon signed-rank tests, two-tailed, Fig 2D). Thus, CS choices during choice-induced revaluation increased, whereas non-choices decreased our measure of hippocampal CS-US associative strength. However, while there was evidence for significant PRE-POST reduction in repetition suppression for CS^0^_A_ (*Z* = 1.84, *P* = .033, *U*_*3*_ = .69, Wilcoxon signed-rank test, one-tailed), the PRE-POST increase in repetition suppression for CS^+^_A_ was not significant (*Z* = 1.48, *P* = .070, *U*_*3*_ = .62, Wilcoxon signed-rank test, one-tailed).

**Fig. 2.**
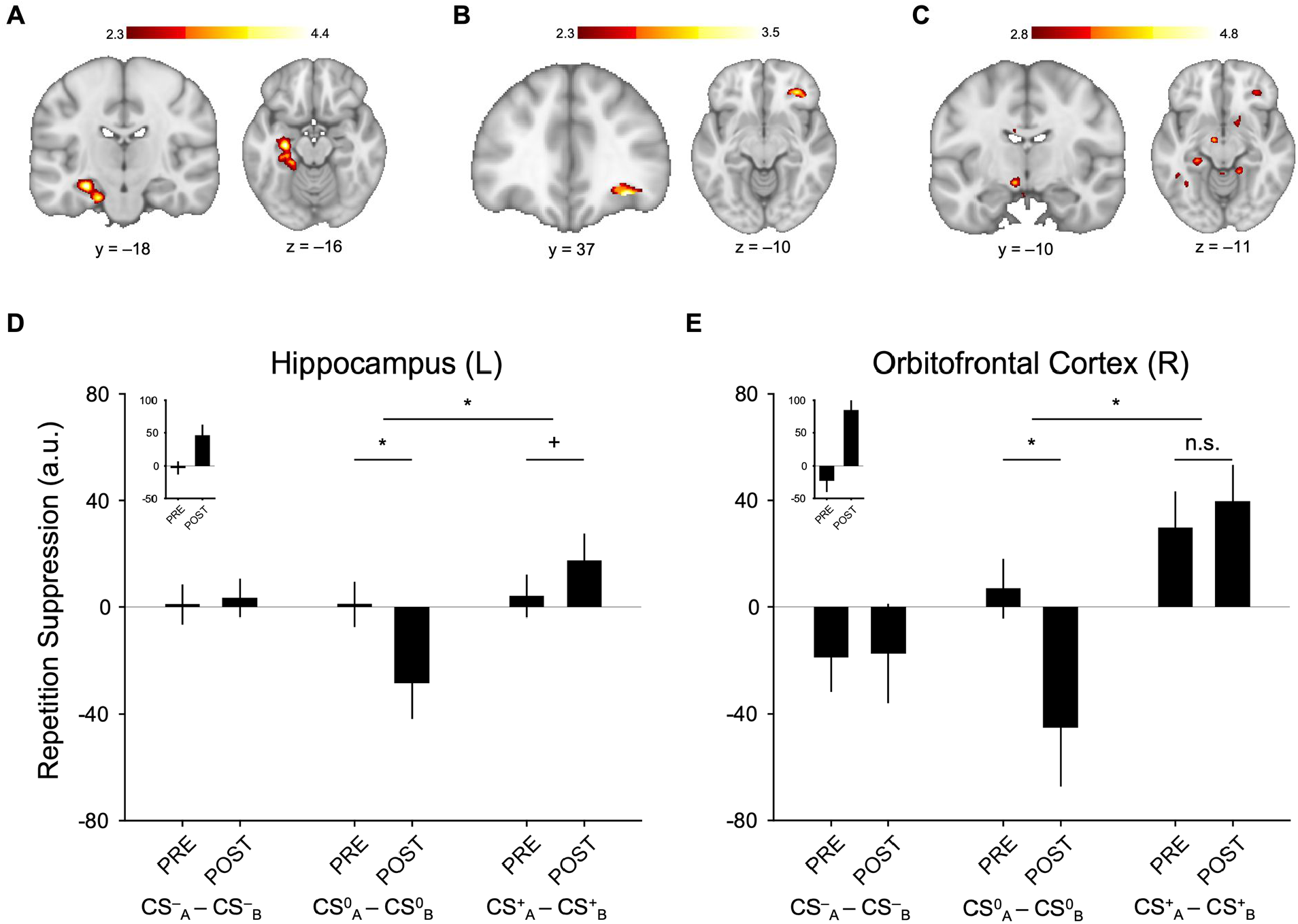
Whole-brain effects of choice-induced revaluation. A) Left hippocampus encodes conjunction of decreased repetition suppression for CS^0^_A_-US^0^ relative to CS^0^_B_-US^0^, controlling for activation elicited by CS^0^_A_ and CS^0^_B_ followed by both incorrect outcomes (US^-^ and US^+^) (Equation 1) AND increased CS^+^_A_-US^+^ repetition suppression relative to CS^+^_B_-US^+^, controlling for activation elicited by CS^+^_A_ and CS^+^_B_ followed by both incorrect outcomes (US^-^ and US^0^) (Equation 2). B) Using small-volume correction (independent mask from Ref. (Jocham et al., 2016)), the same effect was found in the right lateral orbitofrontal cortex (lOFC). C) At a lenient threshold of *Z* > 2.8 (uncorrected) we observed the same effect in left ventral tegmental area and right nucleus accumbens. D) Extracted parameter estimates of the effect in left hippocampus. E) Extracted parameter estimates of the effect in right lOFC.

Furthermore, we found a conjunction effect of a decrease in repetition suppression for CS^0^_A_-US^0^ relative to CS^0^_B_-US^0^ and an increase in repetition suppression for CS^+^_A_-US^+^ relative to CS^+^_B_-US^+^ in the right lateral orbitofrontal cortex that survived small-volume correction (Fig. 2B). Extraction of parameter estimates from this cluster using an independent region of interest (Jocham et al., 2016) revealed that the interaction effect (CS value and time (PRE or POST), *F*_1, 41_ = 5.57, *P* = .023, η^2^_*p*_ = .12, 1–*β* = .99, rmANOVA) was driven by a significant PRE-POST reduction in repetition suppression for CS^0^_A_ (Fig. 2E, *Z* = 1.77, *P* = .039, *U*_*3*_ = .55, Wilcoxon signed-rank test, one-tailed) but not by a PRE-POST increase in repetition suppression for CS^+^_A_ (*Z* = 0.65, *P* = .260, *U*_*3*_ = .50, Wilcoxon signed-rank test, one-tailed). However, we found decreased repetition suppression for CS^0^_A_ (*Z* = 1.74, *P* = .040, *U3*_*1*_ = .55, Wilcoxon signed-rank test, one-tailed) and increased repetition suppression for CS^+^_A_ (*Z* = 2.68, *P* = .004, *U3*_*1*_ = .67, Wilcoxon signed-rank test, one-tailed) during POST, without evidence for non-zero differences in PRE (all *Zs* < 1.77, *Ps* > .077, *U3*_*1*_ < .62, Wilcoxon signed-rank tests, two-tailed). Exploratory analyses at a lenient, uncorrected threshold (*Z* > 2.8, uncorrected) yielded clusters in the left ventral tegmental area (VTA) and the right nucleus accumbens (NAcc, Fig 2C). Both effects were driven by significant reductions in repetition suppression for CS^0^_A_ (VTA: *Z* = 1.81, *P* = .035, *U*_*3*_ = .79; NAcc: *Z* = 2.38, *P* = .009, *U*_*3*_ = .64, Wilcoxon signed-rank tests, one-tailed), but showed only NAcc evidence of increases in repetition suppression for CS^+^_A_ (VTA: *Z* = 0.41, *P* = .340, *U*_*3*_ = .55; NAcc: *Z* = 1.53, *P* = .064, *U*_*3*_ = .64, Wilcoxon signed-rank tests, one-tailed). Overall, these repetition suppression results suggest that decisions during choice-induced revaluation had clearly dissociable effects on the neural representation of previously learned CS-US associations: While the previously chosen CS exhibited increased associative strength to its related US, the exact opposite effect was true for the previously unchosen CS, exhibiting decreased associative strength to its related US. Importantly, the observed dissociation of choice-induced increase of RS effects for CS^+^_A_ choice-induced decrease of RS effects for CS^0^_A_ and the absence of PRE-POST differences of RS effects for the CS^−^_A_ relative to CS^−^_B_ cannot be readily explained by general extinction effects resulting from exposition to CS-US associations other than the initially learned associations. Extinction would have been expected to equally spread across all CS-US associations and would imply equidirectional changes of all CS-US associations from PRE to POST, which is incompatible with the observed results.

### Choice-induced decrease of multivariate neural pattern similarity

Complementary to the mass-univariate repetition suppression-based approach, we performed multivariate fMRI analyses, employing a neural pattern similarity analysis (a variant of representational similarity analysis, RSA (Kriegeskorte et al., 2008)) in the left hippocampus and right lateral OFC. Using the same logic as for the repetition suppression-based analyses, we reasoned that presentation of a CS would induce pre-activation of neural ensembles representing the associated US. This mnemonic pre-activation should not only be present in trials where the CS was followed by the originally learned US, but also in trials where the CS was followed by any of the other two possible, but not associatively linked US. Similarity of neural patterns related to two CS from the same value category could thus be indicative of associative memory retrieval of a US representation. According to the idea of choice-induced weakening of CS^0^_A_ association with US^0^ and strengthening of CS^+^_A_ association with US^+^, our hypothesis therefore was that neural similarity between same value stimulus-outcome pairs (CS^0^_A_-US^−^/CS^0^_B_-US^−^ and CS^0^_A_-US^+^/CS^0^_B_-US^+^; CS^+^_A_-US^−^/CS^+^_B_-US^−^ and CS^+^_A_-US^0^/CS^+^_B_-US^0^) should decrease from PRE to POST, indicating less similarity between patterns of interest (i.e. the weakened/strengthened stimulus and the respective partner stimulus). Therefore, we used one-tailed tests accordingly. For the pair of control stimuli (CS^-^_A_-US^0^/CS^-^_B_-US^0^ and CS^-^_A_-US^+^/CS^-^_B_-US), we did not expect changes in neural pattern similarity and thus employed two-tailed tests.

In the left hippocampal ROI, we observed significant negative PRE-POST change in neural pattern similarity when averaging across all patterns of interest (*t*_41_ = 2.09, *P* = .021, *U3*_*1*_ = .64, 1–*β* = .63, one-sample t-test, one-tailed) and for the CS^+^_A_/CS^+^_B_ pairs (*t*_41_ = 1.81, *P* = .039, *U3*_*1*_ = .57, 1–*β* = .53, one-sample t-test, one-tailed), but only a numerical decrease of neural pattern similarity from PRE to POST for the CS^0^_A_ -/CS^0^_B_ pairs (*t*_41_ = 1.01, *P* = .144, *U3*_*1*_ = .50, 1–*β* = .28, one-sample t-test, one-tailed). Importantly, change of neural pattern similarity for the control stimulus pairs CS^−^_A_/CS^−^_B_ was numerically positive, indicating descriptively higher similarity of CS^0^_A_-/CS^0^_B_ across time, and there was also no evidence of change in neural pattern similarity for the control stimulus pairs CS^−^_A_ /CS^−^_B_ (*t*_41_ = 0.76, *P* = .451, *U3*_*1*_ = .57, 1–*β* = .12, one-sample t-test, two-tailed, Figure 3A).

**Fig. 3.**
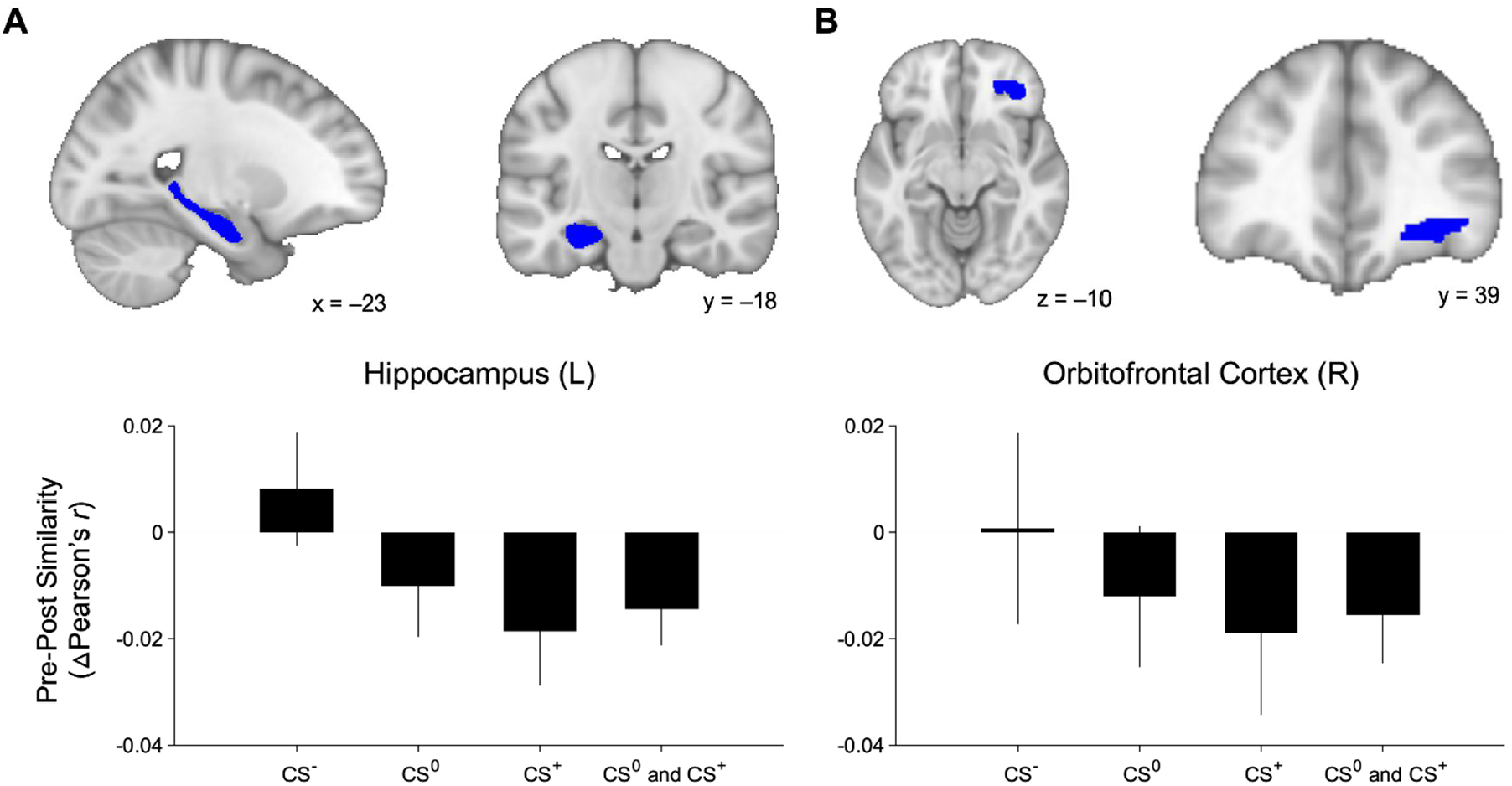
Multivariate Neural Pattern Similarity Analysis Results. A) Multivariate neural pattern similarity analyses in the left hippocampus and B) right lateral OFC, showing that neural similarity (POST *r* – PRE *r*, Δ Pearson’s *r*) between same value stimulus-outcome pairs (CS^0^: CS0A-US^−^/CS^0^_B_-US^−^ and CS^0^_A_-US^+^/CS^0^_B_-US^+^; CS^+^: CS^+^_A_-US^−^/CS^+^_B_-US^−^ and CS^+^_A_-US^0^/CS^+^_B_-US^0^) decreases from PRE to POST. Importantly, change of neural pattern similarity for the control stimulus pairs CS^−^_A_/CS^−^_B_ (CS^−^) is numerically positive, indicating higher similarity from PRE to POST.

In the right lOFC ROI, we observed qualitatively similar results as in the left hippocampus: There was significant negative PRE-POST change in neural pattern similarity when averaging across all patterns of interest (*t*_41_ = 1.70, *P* = .049, *U3*_*1*_ = .62, 1–*β* = .40, one-sample t-test, one-tailed). However, unlike in the hippocampus there was no evidence of significant change in pattern similarity for the CS^+^_A_/CS^+^_B_ pairs (*t*_41_= 1.23, *P* = .113, *U3*_*1*_ = .64, 1–*β* = .23, one-sample t-test, one-tailed). There was also no evidence for a decrease of neural pattern similarity from PRE to POST for the CS^0^_A_ -/CS^0^_B_ pairs (*t*_41_ = 1.55, *P* = .061, *U3*_*1*_ = .62, 1–*β* = .13, one-sample t-test, one-tailed). Only numerically, both CS^+^_A_/CS^+^_B_ pairs and CS^0^_A_ -/CS^0^_B_ pairs became less similar from PRE to POST. The change of neural pattern similarity for the control stimulus pairs CS^−^_A_/CS^−^_B_ was positive, but not significantly different from 0 (*t*_41_ = 0.04, *P* = .969, *U3*_*1*_ = .43, 1– *β* = .05, one-sample t-test, two-tailed, Figure 3B).

Taken together, these multivariate pattern similarity results conceptually confirm the main finding from the mass-univariate repetition suppression-based analyses and further support the interpretation that the observed choice effects could be explained by choice-induced changes of associative strength. However, these results should be interpreted with caution, as power was generally low (1–*β* < .80), most likely resulting from the reduced number of trials included in the analyses (40 trials per CS pair in PRE and POST). Additionally, unlike our repetition suppression-based results, changes in neural pattern similarity do not allow to infer the directionality of the effects (i.e. patterns may become more dissimilar both due to strengthening or weakening of the associative trace). Nevertheless, the results from both sets of analyses provide convergent evidence for our hypothesis that revaluation choices changed the degree to which neural US representations were pre-activated by their associated CS.

### Hippocampal CS-US repetition suppression correlates with future choices

We next investigated whether the observed choice-induced modifications of hippocampal CS-US repetition suppression were correlated with choice biases during the probe phase. As in Experiment 1 and 2, we had formulated directional hypotheses for the choice effects (increased choice probabilities for the previously chosen and decreased choice probabilities for the previously unchosen CS), and thus used one-tailed post-hoc tests.

Unlike in Experiment 1, CS^0^_A_ (unchosen stimulus during revaluation) was not chosen at a lower rate than CS^0^_B_ in the probe phase (*Z* = 1.01, *P* = .844, *U*_*3*_ = .55, Wilcoxon signed-rank test, one-tailed, Fig. 1H) in Experiment 4. Relatedly, there was no evidence for preference differences in within-category choice trials directly comparing CS^0^_A_ and CS^0^_B_ (*Z* = 1.07, *P* = .857, *U3*_*1*_ = .62, one-sample Wilcoxon signed-rank test vs. 0.5, one-tailed; Supplementary Fig. S2H). To assess PRE to POST changes of associative strength participants in the present study had to be re-exposed with the initially learned CS-US associations and were explicitly instructed to judge whether the presented CS-US associations were correct. It is well established that restudying of memorized material reverses retrieval-induced forgetting effects (Hulbert & Norman, 2015; Storm, Bjork, & Bjork, 2008). Thus, the observed behavioral null effect for CS^0^_A_ might be due to re-exposure and restudy of the original CS-US association. This process might have allowed the weakened association between CS^0^_A_ -US^0^ to regain its original associative strength.

However, replicating Experiment 1, we observed a choice-induced increase in preference for CS^+^_A_ over CS^+^_B_ (*Z* = 3.03, *P* = .001, *U*_*3*_ = .76, Wilcoxon signed-rank test, one-tailed, Fig. 1H). This effect was mainly driven by preference for CS^+^_A_ in within-category choice trials directly contrasting CS^+^_A_ and CS^+^_B_ (*Z* = 1.93, *P* = .027, *U3*_*1*_ = .62, one-sample Wilcoxon signed-rank test vs. 0.5, one-tailed; Supplementary Fig. S2H). We therefore focused on this effect in brain-behavior correlations. We hypothesized a positive linear relationship between the difference between choice probabilities of CS^+^_A_ and CS^+^_B_ and the magnitude of hippocampal repetition suppression between CS^+^_A_-US^+^ and CS^+^_B_-US^+^), and thus used one-tailed tests on the Spearman correlation coefficient. The difference of hippocampal repetition suppression between CS^+^_A_-US^+^ and CS^+^_B_-US^+^ was positively correlated with the difference between choice probabilities of CS^+^_A_ and CS^+^_B_ (ρ_41_ = .31, *P* = .024, Spearman correlation, one-tailed; Fig. 4B). The more the hippocampal representation of the CS^+^_A_-US^+^ association had been strengthened by choices during revaluation, the more likely participants were to select CS^+^_A_ compared to its non-revalued partner stimulus CS^+^_B_.

**Fig. 4.**
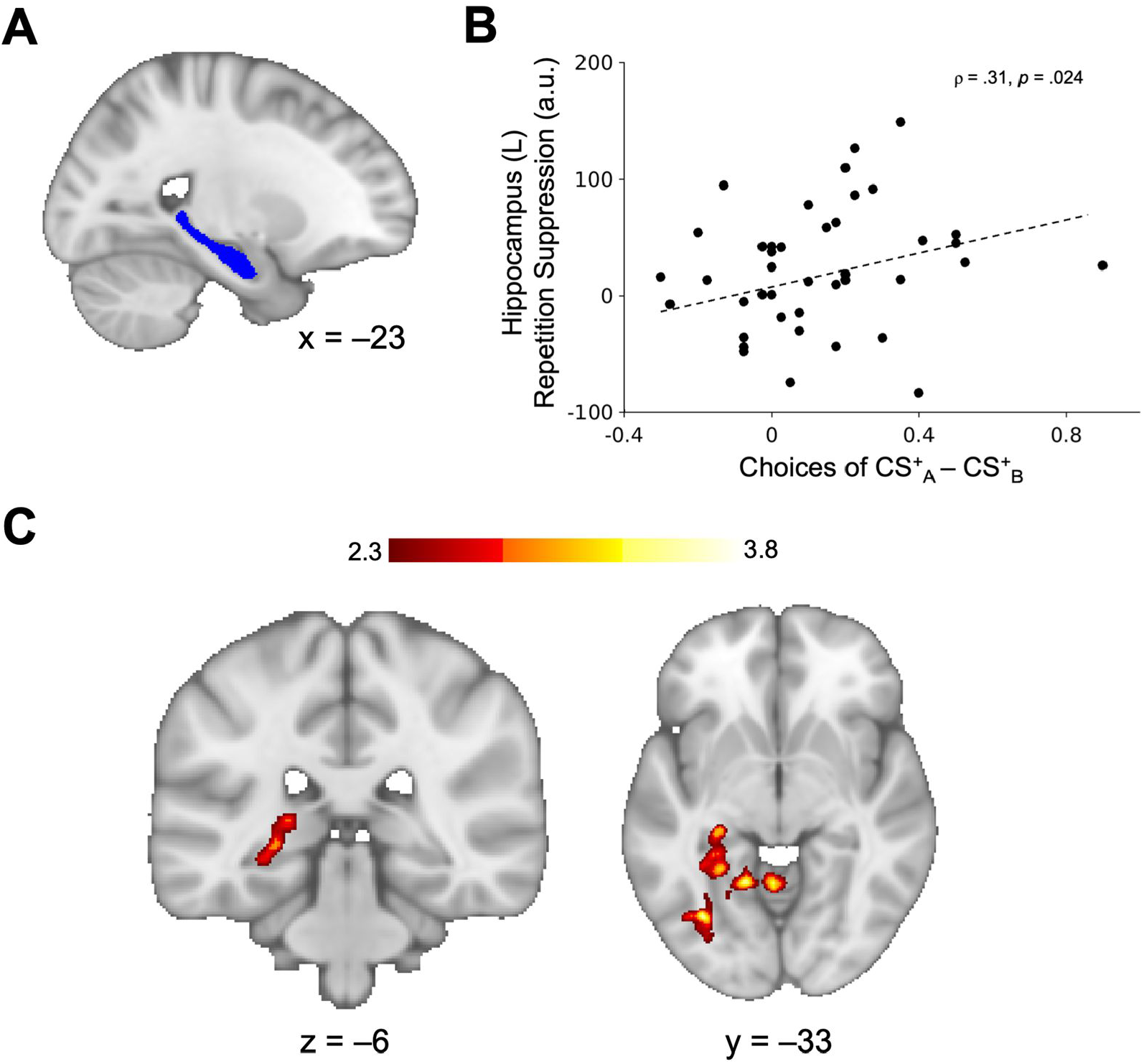
Brain-behavior correlation and whole-brain regressions. B) Left hippocampus repetition suppression of CS^+^_A_-US^+^ relative to CS^+^_B_-US^+^, controlling for activation elicited by CS^+^_A_ and CS^+^_B_ followed by both incorrect outcomes (US^-^ and US^0^) (Equation 2), extracted from an independent anatomical mask (A) during POST was positively related to decision probe difference between overall choice probability of CS^+^_A_ and overall choice probability of CS^+^_B_. With increasing repetition suppression, participants were more likely to prefer CS^+^_A_ over CS^+^_B_. C) Whole-brain regression showing a positive relationship between the difference of CS^+^_A_ and CS^+^_B_ preference and PRE-POST change of repetition suppression in the left posterior hippocampus. With more positive PRE-POST repetition suppression change, participants are more likely to prefer CS^+^_A_ over CS^+^_B_.

In line with this, in a whole-brain analysis we observed a positive relationship between the difference in choice preference for CS^+^_A_ versus CS^+^_B_ and changes in CS-US repetition suppression from PRE to POST in the left posterior hippocampus, extending to left occipito-temporal complex (Fig. 4C). A similar whole-brain analysis using only the choices between CS^+^_A_ and CS^+^_B_ as behavioral covariate revealed areas in the bilateral anterior insula and orbitofrontal cortex (Supplementary Fig. S4B). Neither VTA, NAcc, nor the cluster in the lateral orbitofrontal cortex showed relationships with probe phase behavior.

As an alternative to the proposed associative mechanism, parts of our results could also be explained based on cached values. During Pavlovian conditioning, participants might acquire incentive (“cached”) CS values and use these to guide their decisions, independent of CS-US associations. It is possible that revaluation choices changed these cached CS values, instead of the CS-US associations. Under this reasoning, one would assume that the value information conveyed by CS^0^_A_, the CS not chosen during revaluation and supposedly devalued, would become more similar to the value information of the low-valued US^−^ after choice-induced revaluation (Summerfield, Luyckx, & Sheahan, 2019). By its very nature, the fMRI-RS signal for learned CS-US associations represents BOLD signal reductions due to repetition of both value and identity features shared by CS and US, alongside the associative strength between CS and US (Klein-Flugge et al., 2013). However, CS^0^_A_ should by design of the experiment not be capable to elicit any associative strength- or identity-related RS effects when followed by US^−^, as it was never coupled with US^−^ during Pavlovian conditioning. Since value and identity of the US are inextricably linked in the present design, we reasoned that any RS signal changes for CS^0^_A_ followed by US^−^ from PRE to POST would most likely be attributable to changes in valuation of CS^0^_A_. The cached value account would predict larger repetition suppression effects for CS^0^_A_ followed by US^−^ as for CS^0^_B_ followed by US^−^ during POST. Our design allowed us to set up a contrast to test this possibility (Methods – fMRI contrasts). This indeed yielded effects in the right posterior hippocampus and a midbrain region in the vicinity of the dorsolateral substantia nigra pars compacta (Supplementary Fig. S3) in POST. However, consistent with the absence of a behavioral revaluation effect for CS^0^_A_, the observed hippocampal effect did not differ PRE-POST (test on parameter estimates extracted from right hippocampal anatomical mask, *Z* = 0.90, *P* = .184, *U*_*3*_ = .64, Wilcoxon signed-rank test, one-tailed). Importantly, the parameter estimates of the “cached value” effect in the right hippocampus and the “associative” effect in the left hippocampus were not correlated (all ρ*s* < .18, *Ps* > .260, Spearman correlations, two-tailed). Even when extracting parameter estimates for “cached value” (Equation 4) and “associative” effect (Equation 1) from the exact same anatomical mask of the left hippocampus, we did not observe significant correlations between the two contrasts (PRE: ρ = .22, *P* = .170; POST: ρ = –.03, *P* = .852; Spearman correlations, two-tailed), which would have been expected if both contrasts measure the same with flipped signs. According to the cached value account, hippocampal representations of CS^0^_A_ -US^−^ should be inversely related to preferences of CS^0^_A_. However, directly opposing this prediction, relationships of right hippocampus parameter estimates, and choice behavior were positive (all ρ*s* < .31, *P* > 0.05, Spearman correlations, two-tailed), rendering an explanation of the observed behavioral results based on cached values unlikely. It should be noted that the “cached value” effects would have critically depended on choice-induced devaluation of CS^0^_A_. However, as we did not find behavioral support for the hypothesized devaluation effect, the “cached value” effects should be interpreted with caution.

## Discussion

Using a novel paradigm, we showed that decisions bias future choices, even without participants directly experiencing the outcomes of their decisions. Participants were more likely to select CS they had previously chosen, and less likely to select CS they had not chosen previously, compared to otherwise equivalent CS. At the neural level, we found that choices introduced alterations to hippocampal and orbitofrontal representations of stimulus-outcome associations that were correlated with future decisions.

The idea that past decisions bias preferences was put forth decades ago (Ariely & Norton, 2008; Brehm, 1956) and compelling evidence for post-decision revaluation has accumulated since (Ariely & Norton, 2008; Izuma et al., 2010; Riefer et al., 2017; Sharot et al., 2010) (but see Chen & Risen, 2010 for critical discussion). Here, we present behavioral evidence from four independent data sets that reward-predictive CS that were chosen in the past are more likely to be selected during future choice contexts, compared to CS of equal value that were not presented during choice-induced revaluation. Conversely, we found a decreased preference for CS that were not chosen in the past, compared to otherwise equivalent CS, indicating dissociable effects of choice-induced revaluation. Most importantly, our behavioral findings are independent of experienced rewards, as participants were never presented with the outcomes of their choices. This suggests that the observed effect could arise from associative memory mechanisms, as also indicated by an additional control experiment which was specifically designed to rule out the alternative explanation that the observed choice biases could result from learned choice heuristics. Computational modelling further favored an associative process that differentially updated CS-US associative strength of chosen and unchosen options based on “fictive” reward prediction errors elicited by associative retrieval during decisions in the choice-induced revaluation phase.

Our results are in line with previous reports of choice-induced preference changes (Ariely & Norton, 2008; Brehm, 1956; Izuma et al., 2010; Sharot et al., 2010) and conceptually replicate studies showing changes in stimulus valuation by cued approach training (CAT) (Aridan, Pelletier, Fellows, & Schonberg, 2019; Botvinik-Nezer, Bakkour, Salomon, Shohamy, & Schonberg, 2019; Salomon, Botvinik-Nezer, Oren, & Schonberg, 2019; Schonberg et al., 2014). Similar to the present approach, performance of a button press (“go” response) upon presentation of “go” stimuli during CAT reliably induces long-term (Salomon et al., 2019) non-reinforced changes of desirability and choice probabilities of “go” stimuli over “no-go” stimuli. CAT effects are independent of initial value of the stimuli and seem to depend on integrity (Aridan et al., 2019) and activation (Salomon et al., 2019; Schonberg et al., 2014) of ventromedial prefrontal cortex, and interactions between orbitofrontal cortex and ventral striatum (Salomon et al., 2019). Importantly, the results of Experiment 5 suggest that choice-induced revaluation effects, at least for previously chosen options, seem to go beyond a trained action tendency or choice rule (“go” response), as observed in CAT.

As most previous studies have presented participants directly with the outcomes to be chosen and thereby confounded individual contributions of memory and choice mechanisms, our study might be the first to provide evidence for a value-independent associative memory mechanism driving choice-induced preference changes. Furthermore, even though associative memory dynamics appear as a likely candidate mechanism for choice-induced preference changes, the exact neural mechanisms underlying this decision making phenomenon have remained unknown. Here, we provide evidence that past choices bias future decision making, in part by modifying the strength of neural stimulus-outcome associations. The present results suggest that, during decision making, reactivation of stimulus-outcome associations (Barron et al., 2013; Boorman et al., 2016; Klein-Flugge et al., 2013; Wimmer & Shohamy, 2012) and making a choice renders them subject to nonmonotonic plasticity, with the association of the chosen stimulus being strengthened, and the association of the unchosen stimulus being diminished. Since both chosen and unchosen CS activate neural populations representing the respective associated outcome (Barron et al., 2013; Boorman et al., 2016; Howard et al., 2016; Klein-Flugge et al., 2013; Onat & Büchel, 2015; Tonegawa et al., 2018), we reason that, consistent with the Nonmonotonic Plasticity Hypothesis (reviewed in Ritvo et al., 2019), the observed opposing decision biases are presumably related to additional choice-induced activation of the chosen CS-US association, and absence of such choice-induced activation of the unchosen CS-US associations. However, it should be noted that we did not observe behavioral evidence for choice-induced weakening of the unchosen CS-US association in Experiment 4. This might be due to re-exposure to the initially learned CS-US associations during the POST fMRI run, which presumably allowed the weakened association between CS^0^_A_ -US^0^ to regain its original associative strength, in line with studies showing that restudying of memorized material reverses retrieval-induced forgetting effects (Hulbert & Norman, 2015; Storm et al., 2008). Our results suggest that choices themselves can act as self-generated “teaching signals” (Guggenmos, Wilbertz, Hebart, & Sterzer, 2016; Palminteri et al., 2015), dynamically altering stimulus-outcome associations stored in memory by shifting associative memories along a nonmonotonic plasticity function (Ritvo et al., 2019).

Even though our data provide evidence for choice-induced changes to associative strength, the current approach does not allow to dissociate which exact features of the US contribute to repetition suppression effects. As US value and identity are inextricably linked in our experiment, the observed effects could be related to the quantitative or qualitative changes in CS-dependent pre-activation of identity- or value-related features of the US. However, both identity- and value-related features of the US are learned associatively in the present study and are therefore quite likely to be retrieved in an associative fashion to guide choices. Alternatively, the observed change in the hippocampal repetition suppression could be due to choice-induced alterations of CS representations per se. However, changes of CS representation cannot explain the specificity of RS signals to presentations of the “correct” (learnt) CS-US associations, and, perhaps more importantly, cannot account for the reversed directionality of RS effects depending on choice during revaluation (decrease for CS^0^-US^0^, increase for CS^+^-US^+^), as observed in the data.

Our results suggest that merely making a choice induces plasticity of associative representations in the hippocampus and lateral orbitofrontal cortex. In line with the present results, a fronto-hippocampal network comprising hippocampus and lateral OFC seems to be critically involved in reward-related updating of stimulus-outcome associations (Boorman et al., 2016). Whereas the hippocampus has been shown to be a key brain structure supporting relational learning and memory processes, including value spread during reinforcement learning (Wimmer & Shohamy, 2012), factorized replay of event trajectories (Liu, Dolan, Kurth-Nelson, & Behrens, 2019), the lateral OFC has been additionally implicated in the resolution of credit assignment problems in reinforcement learning (Jocham et al., 2016; Walton, Behrens, Buckley, Rudebeck, & Rushworth, 2010).

Our study has at least three implications for current theories of decision making. First, it proposes a memory-based account for choice-induced revaluation and, more broadly, choice history bias, two well-known, but still poorly understood decision making phenomena. By teasing apart decisions and memory processes related to choice-induced revaluation, we show that choices are not only guided by associative representations of value stored in memory, but that decisions themselves dynamically transform associative memories. This suggests that relational structures stored in decision makers’ cognitive maps (Behrens et al., 2018; Tolman, 1948) can be distorted through their very own choice behavior. Second, we provide evidence for involvement of the hippocampus and lateral OFC in maintaining and updating of stimulus-outcome associations. This extends previous findings (Boorman et al., 2016; Jocham et al., 2016; Klein-Flugge et al., 2013; Walton et al., 2010) by describing a functional role of the hippocampus in value-based decision making that is independent of experienced reward. Such a role may more closely resemble naturalistic decision situations where consequences of choices often unravel at distant future time points, rendering credit assignment challenging (Jocham et al., 2016; Walton et al., 2010). Third, we provide a mechanism underlying seemingly irrational choice behavior: Even though participants chose between equivalent options, choice-induced revaluation biased them towards preferring chosen, and towards neglecting non-selected options. The latter might have important implications for explaining subjective preferences, especially in consumer choice behavior (Riefer et al., 2017) and in understanding why humans tend to make coherent decisions, even in conditions characterized by maladaptive choice behavior such as substance dependence or obsessive compulsive disorder.

Taken together, both our behavioral and neural results support the key prediction that past choices bias future decision making, partially by altering hippocampal and orbitofrontal representations of stimulus-outcome associations. Our study provides a memory-based mechanism to account for choice-induced preferences change effects (Ariely & Norton, 2008; Brehm, 1956; Izuma et al., 2010; Sharot et al., 2010). The present study shows that merely retrieving stimulus-outcome associations and making a choice is sufficient to induce plasticity in reward-predictive associations stored in memory and further suggests that relational structures constituting a decision makers’ cognitive maps can be adaptively altered through their very own choice behavior.

## Methods

### Participants

Participants were recruited from the local student community of the Otto von Guericke University Magdeburg and the Heinrich Heine University Düsseldorf, Germany by public advertisements and via online announcements. Only participants indicating no history of psychiatric or neurological disorder and no regular intake of medication known to interact with the central nervous system were included. Participants in all experiments had normal or corrected-to-normal vision and did not report experience with Japanese kanjis or Chinese characters. All participants provided informed written consent before participation and received monetary compensation for taking part in the study. The study was approved by the local ethics committee at the medical faculty of the Otto von Guericke University Magdeburg, Germany (February 2^nd^ 2018, reference number: 19/18) and conducted in accordance with the Declaration of Helsinki.

49 young, healthy volunteers (age: *M* = 23.93, *SD* = 2.90 years, 18 males) participated in Experiment 1. Seven participants were excluded from statistical analyses due to lacking engagement in the cover task during the Pavlovian conditioning phase that served as an attentional control (<10 % responses in trials that required to indicate the color of the square surrounding the conditioned stimuli. Two additional participants had to be excluded due to not passing the manipulation check (high-value option selected < 50 % during choice-induced revaluation), thus leaving a total of *n* = 40 participants for final analyses.

64 young, healthy volunteers (age: *M* = 23.47, *SD* = 3.79 years, 26 males), participated in experiment 2. Ten participants were excluded from statistical analyses due to lacking engagement in the cover task during the Pavlovian conditioning phase, and thirteen subjects were excluded due to not passing the manipulation check (intermediate valued option selected < 50 % during choice-induced revaluation), one additional participant had to be excluded due to a technical error, leaving *n* = 40 participants for statistical analyses.

61 young, healthy volunteers (age: *M* = 23.26, *SD* = 3.27 years, 23 males), participated in Experiment 3. Ten participants were excluded from statistical analyses due to lacking engagement in the cover task during the Pavlovian conditioning phase, and six subjects were excluded due to not passing the manipulation check (high-value option selected < 50 % during choice-induced revaluation), one additional participant had to be excluded due to a technical error (no data was recorded), leaving *n* = 44 participants for statistical analyses.

58 young, healthy and magnetic resonance imaging (MRI)-compatible volunteers (age: *M* = 24.61, *SD* = 4.01 years, 30 males) participated in Experiment 4 (functional MRI (fMRI) experiment). One participant fell asleep during the post-revaluation fMRI-repetition suppression (fMRI-RS) run, three participants discontinued the MRI acquisition (one due to claustrophobia, two reported a headache during task performance). Twelve additional subjects were excluded due to not passing the manipulation check (high-value option selected < 50 % during choice-induced revaluation), leaving *n* = 42 participants for statistical analyses.

52 young, healthy volunteers (age: *M* = 22.06, *SD* = 3.69 years, 20 males), participated in Experiment 5. 12 subjects were excluded due to not passing the manipulation check (high-value option selected < 50 % during choice-induced revaluation), leaving *n* = 40 participants for statistical analyses.

### Behavioral task - ratings

Participants received written instructions for the experiment and were instructed once again on the computer screen. The experiment was programmed in MATLAB 2012b (MATLAB and Statistics Toolbox Release 2012b, The MathWorks, Inc., Natick, MA, USA), using Psychophysics Toolbox (Brainard, 1997). Before the task, participants rated twenty-five different sweet and high-caloric food items selected from an online database (Blechert et al., 2014). Subjects were instructed to indicate the subjective desirability of the food items by using the “y” and “m” button on a standard German (QWERTZ) computer keyboard to position a red slider bar on a white visual analog scale (VAS) between 0 (not liked) and 100 (very much liked). The lowest-(Experiment 1: *M* = 17.5, *SD* = 20.51; Experiment 2: *M* = 16.65, *SD* = 16.29; Experiment 3: *M* = 18.59, *SD* = 22.15; Experiment 4: *M* = 20.41, *SD* = 20.44), and highest-rated (Experiment 1: *M* = 96.55, *SD* = 7.10; Experiment 2: *M* = 94.85, *SD* = 9.08; Experiment 3: *M* = 95.21, *SD* = 14.52; Experiment 4: *M* = 97.79, *SD* = 4.96; Experiment 5: *M* = 96.63, *SD* = 6.62) food item as well as a food item rated with the median value of all ratings (Experiment 1: *M* = 57.43, *SD* = 11.55; Experiment 2: *M* = 56.23, *SD* = 9.80; Experiment 3: *M* = 57.14, *SD* = 13.95; Experiment 4: *M* = 60.07, *SD* = 11.71; Experiment 5: *M* = 56.48, *SD* = 9.95) were selected for Pavlovian conditioning. We explicitly decided for clearly differentiable pictures of food items in order to elicit activation of differential neural ensembles coding for those stimuli and to facilitate learning of vivid memories of stimulus-outcome associations. Next, subjects rated twenty (eleven in Experiment 1) Japanese kanjis according to subjective value/liking by using the “y” and “m” button on a standard computer keyboard to position a red slider bar on a white visual analog scale (VAS) between 0 (not liked) and 100 (very much liked). The six kanjis rated closest to 50 (equivalent to “neutral”) were selected and their order was randomized before being associated with the food items in Pavlovian conditioning. The subjective values/liking of the six selected kanjis did not differ significantly from each other (main effect of stimulus: all *F*s < 1.39, *P*s > .220, η^2^_*p*_ s < .03, rmANOVA).

### Behavioral task - Pavlovian conditioning

Participants learned to associate the selected kanjis (conditioned stimuli, CS) with differently valued outcomes (unconditioned stimuli, US) by repeatedly observing one CS (2000 ms), followed by an inter-stimulus interval (1000 ms) marked by a fixation cross, and presentation of one US (2000 ms). Each trial was separated by an inter-trial-interval (ITI) marked by a grey screen. The ITI per trial was drawn from a discretized γ-distribution (shape = 5, scale = 0.9) truncated for an effective range of values between 3000 ms and 8000 ms. Across all CS, CS-US couplings were interleaved with 20 % CS-no-US couplings. This was intended to create an association that would not rapidly extinguish due to extinction effects related to not being presented with the associated US during the subsequent choice phases. Each US was associated with two different CS, resulting in three pairs of differently valued CS: CS^+^_A/B_, CS^0^_A/B_ and CS^−^_A/B_. Participants completed 180 trials (19 of the participants included in Experiment 1 had performed 240 trials) of Pavlovian conditioning, 30 (40, respectively) trials per CS-US association. In Experiment 5, participants performed 160 trials and learned associations between four neutrally rated CS and an intermediate value outcome and a high value outcome (40 trials per CS-US association), resulting in two pairs of differently valued CS: CS^+^_80_, CS^+^_20_ and CS^0^_80_, CS^0^_20_. Importantly, one of the two CS in each pair was followed by the outcome with a probability of 80 % (CS^+^_80_ and CS^0^_80_) while the other CS was followed by the US in 20 % (CS^+^_20_ and CS^0^_20_) of the trials. The participants were presented with a cover story for the experiment: They were told to imagine themselves in preparation for a journey to Japan during which they would need to learn kanjis associated with certain food items to be ready to select their favorite sweets in Japanese shops. As an attentional control, we introduced a simple binary classification task, presenting participants with a red or blue square surrounding the CS. Each CS was equally often presented with a red or blue square and color of the square did not predict US contingency. Subjects were instructed to react as quickly and as correctly as possible by pressing “y” upon seeing a blue square and pressing “m” upon seeing a red square surrounding the CS. Pavlovian conditioning was split up in three blocks of 60 (80, respectively) trials, interleaved with self-paced breaks. In Experiment 4, both ratings and Pavlovian conditioning were performed outside the MRI scanner.

Due to a coding error in the script creating pseudo-randomized reward schedules in Experiment 1, 2, & 3 (which we spotted during setup of Experiment 5), CS were not followed by outcomes in exactly 80% of trials equivalently across all CS. For each experiment, we had set up five different reward schedules, assigning different outcome probabilities to each CS. In Experiment 1, participants received the following outcome probabilities on average: CS^−^_A_ (Probability: *M* = .80, range = .70 – .87), CS^−^B (*M* = .78, range = .73 – .88), CS^0^_A_ (*M* = .78, range = .70 – .87), CS^0^_B_ (*M* = .78, range = .63 – .90), CS^+^_A_ (*M* = .80, range = .70 – .90), CS^+^_B_ (*M* = .86, range = .73 – .93). In Experiment 2, participants received the following outcome probabilities on average: CS^−^_A_ (*M* = .80, range = .73 – .87), CS^−^_B_ (*M* = .76, range = .73 – .83), CS^0^_A_ (*M* = .78, range = .70 – .87), CS^0^_B_ (*M* = .78, range = .63 – .90), CS^+^_A_ (*M* = .80, range = .70 – .90), CS^+^_B_ (*M* = .88, range = .83 – .93). In Experiment 3, participants received the following outcome probabilities on average: CS^−^_A_ (*M* = .80, range = .73 – .87), CS^−^_B_ (*M* = .77, range = .73 – .83), CS^0^_A_ (*M* = .79, range = .70 – .87), CS^0^_B_ (*M* = .77, range = .63 – .90), CS^+^_A_ (*M* = .79, range = .70 – .90), CS^+^_B_ (*M* = .88, range = .83 – .93). It should be noted that these minor differences in outcome probabilities between CS cannot account for the observed choice-induced revaluation effects, as the respective chosen CS on average received less outcomes (CS^+^_A_ in Experiment 1) than or an equal number of outcomes (CS^0^_A_ in Experiment 2) as their same-valued partner stimuli. Contrarily, the respective unchosen CS on average received more outcomes (CS^−^_A_ in Experiment 2) than or an equal number of outcomes (CS^0^_A_ in Experiment 1) as their same-valued partner stimuli. Thus, the outcome probabilities assigned to each CS would have worked against the hypothesized effects. Consistently, there were no significant correlations between the outcome probability during Pavlovian conditioning and decision probe overall or pairwise within-category choice probability (ρ*s* < .29, *P*s > .067, Spearman correlations, two-tailed). Importantly, Experiment 4 was not affected from this error, as reward schedules were created with a different script in which we correctly coded that each CS would be followed by an outcome in 80% of the trials.

### Behavioral task - choice-induced revaluation

After completion of Pavlovian conditioning, participants were presented with repeated choices (28 trials) between a CS^+^_A_ versus a CS^0^_A_ (Experiments 1 & 4), CS^0^_A_ versus a CS^-^_A_ (Experiment 2), a CS^0^ _A_ versus either a CS^-^_A_ (14 trials) or a CS^+^_A_ (14 trials) (Experiment 3), or a CS^+^_80_ versus a CS^0^_80_ and a CS^+^_20_ versus a CS^0^_20_ (Experiments 5), interleaved with lure decisions (28 trials) between four other neutrally rated kanjis that had never been presented during Pavlovian learning and thus were not associated with any of the US. The choice-induced revaluation phase served as the crucial manipulation in all experiments and was systematically varied across studies. Choice probability (CP) for the high-value CS served as a control for learning and as a manipulation check. Only participants selecting the higher valued CS more than 50 % (CP ≥ 0.50) were included in the final analysis, as we reasoned that choice-induced revaluation choices would (1) represent a marker of having learned the true associative values of the CS, (2) be a measure for learning, independent of the actual decision probe phase data (avoiding biased and arbitrary decisions for exclusion of participants), and (3) allow us to exclude decision makers showing random, or arbitrary choice behavior. Choice options were presented for 1500 ms and the chosen option was highlighted by a grey square surrounding the chosen CS. If participants did not respond within the time-window, a time-out message was displayed, and the respective trial was repeated at the end of the choice-induced revaluation phase. Order (left/right) of choice options was counter-balanced to avoid simple response patterns or decision rules (e.g. “always press left”). Participants were instructed to imagine themselves in a Japanese shop, where they would like to buy their favorite food items based on the previously learned kanjis (CS). Participants were told that one of the choice trials would randomly be drawn and their choice would determine which food item (US) they would receive as a bonus upon completion of the experiment. Participants selected choice options by pressing the “y” (left option) or “m” (right option) button (left or right index finger on an MRI-compatible response box in Experiment 4). Importantly, participants were not presented with the US related to their chosen or unchosen CS to dissociate the observed effects from outcome-related relearning of CS-US associations. We assumed that presentation of a CS would pre-activate neural ensembles coding for the associated US. Consequently, we expected that choosing a CS would induce strengthening of the chosen option’s CS-US association, whereas not choosing a CS would weaken the unchosen option’s CS-US association.

### Behavioral task - decision probe

Following choice-induced revaluation, participants were presented with repeated binary choices (120 trials) between all possible CS combinations to assess CS preferences. Every CS combination was presented eight times in pseudo-random order. In Experiment 5, participants made choices between the two same-value pairs of stimuli that were differently strong associated to their respective outcomes (CS^+^_80_ versus CS^+^_20_ and CS^0^_80_ versus CS^0^_20_). Here, every CS pair was presented ten times in pseudo-random order. Choice options were presented for 1500 ms. If participants did not respond within this time-window, a time-out message was displayed, and the respective trial was repeated at the end of the decision probe phase. Participants selected choice options by pressing the “y” (left option) or “m” (right option) button (left or right index finger on MR-compatible response box in Experiment 4). Order (left/right) of choice options was counterbalanced. Participants were instructed that their shopping bag was torn, and they had to return to the shop for buying their favorite food items based on the previously learned kanjis (CS). Again, participants were told that one of the choice trials would randomly be drawn and their choice would determine which food item (US) they would receive as a bonus upon completion of the experiment. Importantly, participants were again not presented with the US related to their chosen or unchosen CS.

### fMRI repetition suppression task (Experiment 4)

After Pavlovian conditioning outside the MRI-scanner, we administered two fMRI-RS blocks, one immediately before (“PRE”) and one immediately after (“POST”) the revaluation phase to assess choice-induced effects of fMRI repetition suppression. Every possible combination of CS and US (eighteen combinations) was presented twenty times each (360 trials in total). In one third of the trials, the originally learned CS-US associations were presented, the remaining two third of trials contained incorrect CS-US associations. In every trial, a CS was presented for 700 ms, followed by an interstimulus interval (fixation cross) for 400 ms and a US for 700 ms. The intertrial interval was drawn from a discretized γ-distribution (shape = 2.01, scale = 1), truncated for an effective range of values between 2000 ms and 6000 ms. Order of trials was pseudo-random, between-trial repetition of CS or US did not occur. Additionally, every batch of 18 trials contained every possible combination of CS-US-association to avoid comparison of temporally distal trials and between-trial biases in repetition suppression introduced by e.g. fluctuations of attention, “novelty” or surprise. During both runs of fMRI repetition suppression, participants performed an attentional control task. After a pseudo-random 20 % of trials, participants were presented with probe trials in which they were asked to indicate whether or not the previously seen CS-US-association matched the true CS-US-association learned during Pavlovian conditioning via button presses with their right and left index fingers on an MRI-compatible response box. Correct responses were rewarded with 0.05 € and incorrect responses or time-out trials (without a response by the participant within 2500 ms after onset of the probe trial) resulted in a 0.05 € penalty which would be summed up as a bonus upon completion of the experiment. On average, participants earned a bonus of 5.93 € (*SD* = 1.05). Performance during the attentional control task was generally high (overall probability of correct answers, excluding time-out trials: *M* = .92, *SD* = .06 (*t*_41_ = 41.54, *P* < .001, one-sample t-test vs. chance level (0.5)), with no evidence for a difference in performance between PRE and POST choice-induced revaluation run (PRE: *M* = .92, *SD* = .074; POST: *M* = .92, *SD* = .072; *t*_41_ = 0.06, *P* = .95, paired-samples t-test).

### fMRI acquisition

Two runs of fMRI were recorded with a 3 Tesla Siemens PRISMA MR-system (Siemens, Erlangen, Germany), using a 64-channel head coil. Blood oxygenation level dependent (BOLD) signals were acquired using a multi-band accelerated T2*-weighted echo-planar imaging (EPI) sequence (multi-band acceleration factor 2, repetition time (TR) = 2000 ms, echo time (TE) = 30 ms, flip angle = 80°, field of view (FoV) = 220 mm, voxel size = 2.2 × 2.2 × 2.2 mm, no gap). Per volume, 66 slices covering the whole brain, tilted by approximately 15° in z-direction relative to the anterior–posterior commissure plane were acquired in interleaved order. The first 5 volumes of the functional imaging time series were automatically discarded to allow for T1 saturation. After each run, a B0 magnitude and phase map was acquired to estimate field maps and B0 field distortion during preprocessing (TR = 660 ms, TE 1 = 4.92 ms, TE 2 = 7.38 ms, flip angle = 60°, FoV = 220 mm). Additionally, before the PRE choice-induced revaluation fMRI-RS run, a high-resolution three-dimensional T1-weighted anatomical map (TR = 2500 ms, TE = 2.82 ms, FoV = 256 mm, flip angle = 7°, voxel size = 1 × 1 × 1 mm, 192 slices) covering the whole brain was obtained using a magnetization-prepared rapid acquisition gradient echo (MPRAGE) sequence. This scan was used as anatomical reference to the EPI data during the registration procedure.

### Behavioral analyses

Data were analyzed using MATLAB 2012b, 2017a and 2019a (The MathWorks, Inc., Natick, MA, USA) using custom analysis scripts. For the manipulation check, indicating learning of the CS-US-associations, average choice probabilities (CPs) for the higher valued CS were calculated (one average in Experiment 1, 2 and 4, two averages in Experiment 3 and Experiment 5) by summing up choices of the higher valued CS and dividing by the number of choice trials with a recorded response. This CP was compared against the inclusion criterion of CP ≥ 0.50. If participants had chosen the high-valued CS in more than 50 % (or in exactly 50 %) of the trials of choice-induced revaluation, they were included in the final analyses. For the decision probe phase, we computed an average overall CP per conditioned stimulus per subject including all binary decisions in which the respective CS was present (count data, 1 representing selection of the CS, 0 representing selection of the alternative CS). Lacking a non-parametric alternative to the parametric two-way repeated-measures analysis of variances (rmANOVA), distributions of mean overall CPs were analyzed at the group level with a rmANOVA with factors CS valence (3) and stimulus type (2) and post-hoc Wilcoxon sign-rank tests for paired samples, focusing on the pairwise comparison of stimulus types within each level of valence. We hypothesized a main effect of CS valence and an interaction effect of CS valence × CS type (A vs. B), resulting from higher CPs for previously chosen CS relative to the equivalent control CS and lower CPs for previously unchosen CS relative to the equivalent control CS (Experiment 1, 2 and 4). However, we expected absence/no evidence for such an interaction effect in Experiment 3 in which a CS^0^_A_ was chosen and unchosen equally often (resulting in no change of the preference relative to the control CS). Additionally, we performed one-sample Wilcoxon sign-rank tests of overall CP against chance level (CP = 0.50). As an alternative measure of choice preference in addition to overall CP, we also computed pairwise choice probabilities by comparing choice ratios of the two CS within each valence category with one-sample Wilcoxon sign-rank against chance level (CP = 0.50).

Experiment 5 was specifically designed to rule out the possibility that participants did not guide their choices based on (altered) associative strength between CS and US but had simply learned choice rules for the two CS presented during the revaluation phase (“go” tag for chosen CS, “no-go” tag for unchosen CS). We thus aimed to orthogonalize contributions of associative strength and choice rule, by assigning “go” tags to the two chosen CS^+^ that differed in their associative strength to US^+^ and “no-go” tags to the two unchosen CS^0^ that differed in their associative strength to US^0^. Our hypothesis was that if choice behavior was exclusively driven by these tags participants had learned “go” tags for both chosen CS^+^_80_ and CS^+^_20_ and “no-go” tags for both unchosen CS^0^_80_ and CS^0^_20_, there should be no evidence for both same-value pairwise choice probabilities different from chance level (CP = 0.5). However, if the choices were made based on the learned associations and the associative strengthening/weakening of the memory trace between CS and US, there should be a significantly increased choice probability for CS^0^_80_/CS^+^_80_ that were more strongly associated with their respective outcomes.

As we had formulated directional hypotheses for the choice effects, we performed one-tailed (post-hoc) tests. In Experiment 3, there were no directional hypotheses for the choice effects, therefore, we used two-tailed post-hoc tests. We report effect sizes η^2^_*p*_ for rmANOVAs, Cohen’s *U*_*3*_ for Wilcoxon signed-rank tests and Cohen’s *U3*_*1*_ for one-sample Wilcoxon signed-rank tests (range: 0 – 1, .5 indicating no effect), calculated in the MATLAB-based Measures-of-Effect-Size-toolbox (Hentschke, 2020). Based on the reported effect sizes η^2^_*p*_ we additionally indicate post-hoc achieved power (1–*β*) for the hypothesized interaction effects of CS valence × CS type in rmANOVAs across behavioral analyses in Experiment 1–4. Based on the means and standard deviations for both choice probabilities, we indicate post-hoc achieved power for Experiment 5. All power analyses were conducted in G*Power (Faul, Erdfelder, Buchner, & Lang, 2009; Faul, Erdfelder, Lang, & Buchner, 2007) (v3.1.9.2).

### Univariate fMRI data analysis

We exploited fMRI repetition suppression effects (rapid, repeated presentation of the same stimulus or pre-activation of a stimulus by associated stimuli elicits reduced neural responses, as stimuli are represented by overlapping neural ensembles (Barron et al., 2016; Grill-Spector & Malach, 2001)) to investigate choice-related changes in neural representations of CS-US associations. As conditioning enhances CS’s ability to pre-activate neural ensembles coding for US, we expected a change of CS-US-associative strength, as measured by repetition suppression after choice-induced revaluation. Strong CS-US associations should elicit high fMRI-RS effects (i.e. low activation), whereas weak CS-US associations should elicit low fMRI-RS effects (i.e. high activation). We expected decreased neural representations of CS-US associations for CS^+^_A_ and increased choice-related neural representations of CS-US associations for CS^0^_A_ after choice-induced revaluation.

All univariate fMRI analyses steps were performed using tools from the Functional Magnetic Resonance Imaging of the Brain (FMRIB) Software Library (FSL, v6.0) (Jenkinson, Beckmann, Behrens, Woolrich, & Smith, 2012). Preprocessing included motion correction using rigid-body registration to the central volume of the functional time series (Jenkinson, Bannister, Brady, & Smith, 2002), correction for geometric distortions using the field maps and an n-dimensional phase-unwrapping algorithm (B_0_ unwarping) (Jenkinson, 2003), slice timing correction using Hanning windowed sinc interpolation, high-pass filtering using a Gaussian-weighted lines filter of 1/100 Hz. EPI images were registered with the high-resolution brain images and normalized into standard (MNI) space using affine registration linear (boundary-based registration) as well as nonlinear registration (Andersson, Jenkinson, & Smith, 2007a, 2007b). Functional data were spatially smoothed using a Gaussian filter with 6 mm full-width at half maximum. We applied a conservative independent components analysis (ICA) to identify and remove obvious artefacts. Independent components were manually classified as signal or noise based on published guidelines (Griffanti et al., 2017), and noise components were removed from the functional time series. General linear models (GLMs) were fitted into prewhitened data space to account for local autocorrelations (Woolrich, Ripley, Brady, & Smith, 2001). The individual level (first level) GLM design matrix per run and participant included fifty box-car regressors in total. Thirty-six regressors coded for onsets and durations of all eighteen presented CS-US-association trials (each modelled as single events of 1800 ms duration), two regressors coded for onsets and durations of the three within-run pauses (each 45 s), two regressors coded for onsets and durations of the attentional control task probe, four regressors coded onsets and durations of left and right button presses (delta stick functions on the recorded time of response button presses) and the six volume-to-volume motion parameters from motion correction during preprocessing were entered. Regressors were convolved with a hemodynamic response function (γ-function, mean lag = 6 s, SD = 3 s). Each first level GLM included five contrasts to estimate individual per run contrasts of parameter estimates for 1) lower CS^0^_A_ -US^0^ repetition suppression relative to the other equivalent CS^0^_B_, controlling for all other combinations of CS^0^ presentations (Equation 1), 2) higher CS^+^_A_-US^+^ repetition suppression relative to the other equivalent CS^+^_B_, controlling for all other combinations of CS^+^ presentations (Equation 2), 3) higher CS^−^_A_ repetition suppression relative to the other equivalent CS^−^_B_, controlling for all other combinations of CS^−^ presentations, 4) Conjunction of 1) & 2), i.e. voxels coding for both decrease of CS^0^_A_ -US^0^ repetition suppression AND increase of CS^+^_A_-US^+^ repetition suppression (Equation 3), 5) right vs. left button press. Two separate pre and post choice-induced revaluation second level (group level) GLMs were carried out by submitting individual level parameter estimates to mixed-effects statistics and ordinary least squares (OLS) regression for higher-level contrast of parameter estimates (COPE) estimation. To control for multiple comparisons, cluster-based correction with an activation threshold of Z > 2.3 using a cluster-extent threshold of *P* < .05 was applied at the whole-brain level.

The key tests for our hypothesis were focused on the effect of revaluation choices on CS-US-associations during the POST run. The PRE run served as a control to rule out potential baseline differences in repetition suppression for CS^0^_A_ or CS^+^_A_.

A priori regions-of-interest (ROIs) comprised the lateral orbitofrontal cortex (lOFC) and the hippocampus, as those regions have been implicated in representation and adaptive changes of stimulus-outcome associations, respectively (Boorman et al., 2016; Garvert et al., 2017; Jocham et al., 2016; Klein-Flugge et al., 2013). We investigated post choice-induced revaluation conjunction effects (Equation 3) to identify regions involved in processing of choice-induced changes to CS-US-associations. An independent functional mask of a contrast investigating stimulus-outcome-associations from a previous study (Jocham et al., 2016) (restricted along the z-direction from –6 to –14 to constrain spatial extent), was used for small-volume correction (*P*_*SVC*_) of the bilateral lOFC. The small-volume corrected functional activation mask from the conjunction contrast was used to extract contrast parameter estimates of the CS^0^ _A_ contrast and the CS^+^_A_ contrast. Additionally, an independent anatomical mask of the hippocampus (Harvard-Oxford Atlas) was used to extract PRE and POST choice-induced revaluation contrast parameter estimates of the CS^0^_A_ contrast and the CS^+^_A_ contrast. PRE versus POST comparisons of activation were carried out using a repeated-measures ANOVA and post-hoc Wilcoxon-sign rank tests. As we had formulated directed hypotheses, and because parameter estimates were extracted from family-wise error corrected ROIs, we used one-tailed post-hoc tests. We report effect sizes η^2^_*p*_ for rmANOVAs, Cohen’s *U3* for Wilcoxon signed-rank tests and Cohen’s *U3*_*1*_ for one-sample Wilcoxon signed-rank tests (range: 0 – 1, .5 indicating no effect), calculated in the MATLAB-based Measures-of-Effect-Size-toolbox (Hentschke, 2020). Based on the reported effect sizes η^2^_*p*_ we additionally indicate post-hoc achieved power (1–*β*) for the hypothesized interaction effects of CS valence × time in rmANOVAs, calculated with G*Power (Faul et al., 2009, 2007) (v3.1.9.2). Additionally, we explored functional activation not surviving whole-brain or small-volume family-wise error corrections by thresholding activation maps at *Z* = 2.8 and extracting contrast estimates of the CS^0^_A_ contrast and the CS^+^_A_ contrast for brain-behavioral correlations.

Extracted parameter estimates were additionally used for brain-behavioral correlations using non-parametric Spearman correlations. For brain-behavior correlations, we had the directed hypotheses that associative strength, as measured by repetition suppression should be positively related to the difference between overall CP of CS^+^_A_ and overall CP CS^+^_B_ and negatively related to difference between overall CP CS^0^_A_ and overall CP CS^0^_B_. Additionally, to refine our insights in the relationship of neural and behavioral results, we also correlated repetition suppression with pairwise CP for CS^+^_A_ versus CS^+^_B_, again assuming a positive relationship and pairwise CP for CS^0^_A_ versus CS^0^_B_, predicting a negative relationship. Due to our directed hypotheses, and because parameter estimates were extracted from family-wise error corrected ROIs, we performed one-tailed tests on Spearman correlation coefficients.

Furthermore, whole-brain voxel-wise regressions were applied at the group level for both PRE and POST run and also a group level analysis on the activation change from PRE to POST run. We used demeaned individual behavioral difference regressors of overall CP of CS^+^_A_ – overall CP CS^+^_B_ and overall CP CS^0^_A_ – overall CP CS^0^_B_ and demeaned pairwise within-category CP of trials directly comparing CS^+^_A_ and CS^+^_B_ and CS^0^_A_ and CS^0^_B_, respectively.

For our proposed associative mechanism, we assumed that after Pavlovian conditioning, a CS would pre-activate the respective, associatively learned US and participants would make their decisions between CS based on the associated outcome. Our hypothesized mechanism relies on the assumption that participants form a (simple) model of the task, which is well in line with the literature on associative learning (Barron et al., 2013; Klein-Flugge et al., 2013). However, alternatively, the observed pattern of results could also be explained by a “cached value” account. According to this account, participants acquire “cached” values during Pavlovian conditioning and further use those “model-free” values to guide their decision, independent of associated outcomes or the learned relationships between CS and US. Our study was not designed to dissociate both mechanisms on a behavioral level. The prediction of the “cached value” account would be that decisions during choice-induced revaluation lead to changes in “cached values” and leave CS-US association unaffected. In other words, CS^0^_A_ should have a reduced “cached value”, whereas the “cached value” of CS^+^_A_ should be increased following choice-induced revaluation. In another fMRI contrast, we tested whether the relationship/similarity between CS^0^_A_ and US^−^ (Equation 4) would change due to the choice-induced revaluation phase. If our results can be explained by the “cached value” account, we would assume that CS^0^_A_ followed by US^−^ would show reduced activation/higher similarity in the post choice-induced revaluation run, compared to its equivalent partner stimulus CS^0^_B_ followed by US^−^. As any other stimulus- or outcome-related effects are controlled for in the contrast, and CS^0^_A_ was not learned to be associated with US^−^ during Pavlovian conditioning (i.e. not being able to pre-activate a US^−^ representation), we assumed that reductions in activation could only be interpreted as a repetition of a shared feature, namely the stimulus/outcome value. After testing the contrast, we extracted parameter estimates from an independent hippocampus mask and correlated the parameter estimates with CP. We hypothesized that CS^0^_A_ and US^−^ similarity would be negatively related to overall CP CS^0^_A_ – overall CP CS^0^_B_. The lower the CS^0^_A_ and US^−^ similarity, the less likely participants would be to select CS^0^_A_.

### Multivariate fMRI data analysis

Complementary to the univariate repetition suppression-based approach, we performed confirmatory multivariate fMRI analyses, employing a neural pattern similarity analysis (a variant of representational similarity analysis, RSA (Kriegeskorte et al., 2008)) in the left hippocampus and right lateral OFC to further support the hypothesized associative strengthening/weakening mechanism. As for the repetition suppression-based analyses we assumed that presentation of a CS should induce pre-activation of neural ensembles coding the respective associated US. This mnemonic pre-activation should not only be present in trials where the CS was followed by the originally learned US, but also in trials where the CS was followed by any of the other two possible, but not associatively linked US. Similarity of neural patterns related to two CS from the same value category could thus potentially be indicative of associative memory retrieval of a US representation. According to the idea of choice-induced weakening of CS^0^_A_ association with US^0^ and strengthening of CS^+^_A_ association with US^+^, our hypothesis was that neural similarity between same value stimulus-outcome pairs (CS^0^_A_ - US^−^/CS^0^_B_ -US^−^ and CS^0^_A_-US^+^/CS^0^_B_ -US^+^; CS^+^_A_-US^−^/CS^+^_B_-US^−^ and CS^+^_A_-US^0^/CS^+^_B_-US^0^) should decrease from PRE to POST, indicating less similarity between patterns of interest (i.e. the weakened/strengthened stimulus and the respective partner stimulus). For the pair of control stimuli (CS^-^_A_-US^0^/CS^-^_B_-US^0^ and CS^-^_A_-US^+^/CS^-^_B_-US), we did not expect changes in neural pattern similarity.

Preprocessing steps for multivariate fMRI analyses were identical as for previously mentioned univariate fMRI analyses, with the only exception that the functional imaging timeseries were not spatially smoothed. As for univariate fMRI analyses, we applied a conservative independent components analysis (ICA) to identify and remove obvious artefacts. General linear models (GLMs) were fitted into prewhitened data space to account for local autocorrelations (Woolrich et al., 2001). The individual level (first level) GLM design matrix per run and participant included fifty box-car regressors in total. Thirty-six regressors coded for onsets and durations of all eighteen presented CS-US-association trials (each modelled as single events of 1800 ms duration), two regressors coded for onsets and durations of the three within-run pauses (each 45 s), two regressors coded for onsets and durations of the attentional control task probe, four regressors coded onsets and durations of left and right button presses (delta stick functions on the recorded time of response button presses) and the six volume-to-volume motion parameters from motion correction during preprocessing were entered. Regressors were convolved with a hemodynamic response function (γ-function, mean lag = 6 s, *SD* = 3 s). Each first level GLM included one contrast to model activation related to each of the eighteen presented CS-US-associations versus baseline (eighteen contrasts in total). The a priori ROIs were built in MNI space and back-projected into subject native space using inverse normalization parameters obtained from FSL during preprocessing procedures. We used these individual ROIs for spatially constrained multivoxel pattern extraction from the respective contrast *t*-value maps. Similarity-based analyses were carried out using the MATLAB-based multivariate pattern analysis toolbox CoSMoMVPA (Oosterhof, Connolly, Haxby, & Rosa, 2016). We employed 1−Pearson’s product-moment correlation coefficient (1−*r*) as a measure of pairwise dissimilarity between neural patterns of interest, separately for PRE and POST and the two ROIs. Within-subject pairwise neural dissimilarity was subtracted from 1 (to create a measure of neural pattern similarity) and Fisher-*Z* transformed to closer approximate normally distributed data. We then calculated the within-subject PRE-POST change between the resulting pairwise neural pattern similarity measures (POST *r* – PRE *r*, Δ Pearson’s *r*). As we had a directional hypothesis of negative PRE-POST change of both CS^0^_A_/CS^0^_B_ and CS^+^_A_/CS^+^_B_ and to increase the number of trials included in the inference, we pooled neural pattern similarity measures across all patterns of interest. Lastly, average neural similarity changes were analyzed at the group level with one-sample t-tests against 0. Due to the expected negative effects of neural pattern similarity changes in patterns of interest, we used one-tailed tests accordingly. As there was no such directional hypothesis for the pairs of CS^−^_A_/CS^−^_B_, we employed two-tailed tests. We report effect sizes Cohen’s *U3*_*1*_ for one-sample t-tests against 0 (range: 0 – 1, .5 indicating no effect), calculated in the MATLAB-based Measures-of-Effect-Size-toolbox (Hentschke, 2020). Based on the mean and standard deviation of pattern similarity measures, we report post-hoc achieved power of all analyses.

### Univariate fMRI contrasts

CS^0^_A_ fMRI-RS contrast:

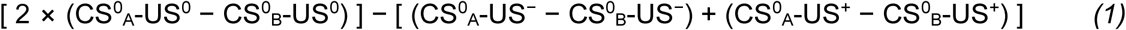

CS^+^_A_ fMRI-RS contrast:

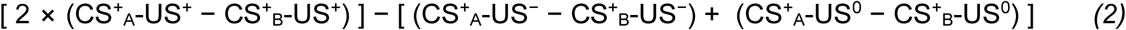

Conjunction fMRI-RS contrast:

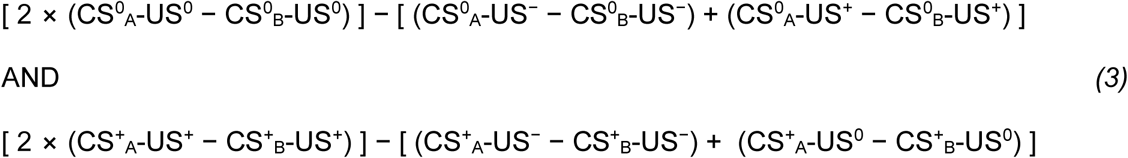

“Cached” value control analysis fMRI-RS contrast:

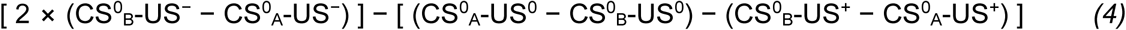

### Computational models

To formally characterize behavior in our experiments, we fit six different variants of a reinforcement learning model using Rescorla-Wagner-like delta update rules (Rescorla & Wagner, 1972). For each experiment, we compared the six models that implemented different ways by which participants could have learned CS-US associative strength - and updated associative strength during choice-induced revaluation.

#### Model 1: Pavlovian learning only

In this model, CS values are exclusively acquired during Pavlovian learning, without any further update during the choice-induced revaluation. On each trial, the value of the stimulus currently presented was updated according to the following rule:

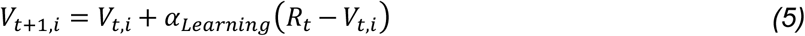

where *α*_*Learning*_ is the learning rate, *V*_*t*_,_i_ is the value of the i^th^ stimulus (1 to 6 for the six CS), and *R*_*t*_ is the reward value of the US (0, 0.5 and 1 for low-, intermediate, and high-value outcome, respectively, and 0 if no outcome was presented) in the Pavlovian conditioning phase. CS values were initialized at 0.5. The estimated associative strength for each CS after the learning phase was directly passed to a softmax choice rule (Equation 7), without further modulation of CS associative strength by choice-induced revaluation.

#### Model 2: Pavlovian learning and choice-induced revaluation (chosen CS)

Model 2 acquired stimulus values during Pavlovian learning exactly like model 1, but additionally updated CS-US associative strength by “fictive reward prediction errors” elicited by decisions in the choice-induced revaluation phase. The “fictive reward prediction errors” was based on our reasoning that presentation of CS during revaluation would lead to retrieval of the associated US and that, consistent with our hypothesis, stimulus-outcome association of the chosen CS would be strengthened, whereas the association of the unchosen CS would be weakened, resulting from nonmonotonic memory plasticity (Ritvo et al., 2019). As no objective feedback (US) was presented during choice-induced revaluation, the reward prediction could only be derived by assuming associative retrieval. This model only updated associative strength of the chosen, but not the unchosen CS.

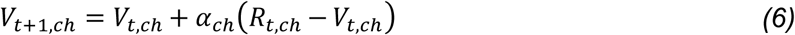

where *α*_ch_ is the choice revaluation learning rate scaling the impact of “fictive” reward prediction errors elicited by fictive outcomes *R*_*t*, ch_ (1 for chosen CS and −1 for unchosen CS) elicited by the US associated with each CS.

#### Model 3: Pavlovian learning and choice-induced revaluation (chosen CS)

This model was set up to account for the possibility that updating of both the chosen and unchosen CS association could occur during the choice-induced revaluation. It was identical to model 2, with the exception that it performed an update to the associative strength of the unchosen stimulus, using a separate learning rate *α*_unch_, in addition to updating the chosen CS associative strength.

These same three models were set up as variants (“associative value models”) that were identical in all respects except for (1) the outcomes during Pavlovian learning, which were modelled as 0 (no outcome presented) or 1 (outcome presented) and CS associative strength values were scaled with the normalized (0-1), individually rated subjective value of the outcome (pre-rating) at the end of the Pavlovian learning phase.

The estimated associative strengths for all CS after the choice-induced revaluation phase were passed to a softmax decision function to generate choice probabilities for each option on each trial:

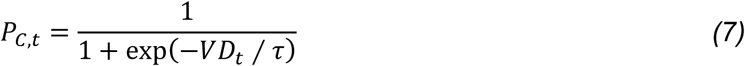

where *P*_*C*_,_*t*_ is the model’s probability to select option C on trial t, the choice the participant actually made on trial t. *VD*_*t*,_ is the value difference (or difference in associative strength) between the chosen and unchosen CS on trial t, and *τ* is a temperature parameter that determines the degree of stochasticity in participants’ choice behavior.

To find the free parameters that best described participants’ behavior, we used a two-step fitting procedure to minimize the negative log likelihood estimate (−LLE):

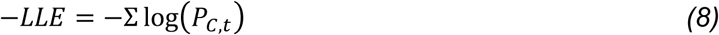

First, we ran a grid search on an n-dimensional grid in log space (where n = number of free parameters for each model), with 30 steps along each dimension. The grid optimum was then used as initial value and passed to constrained non-linear optimization using the MATLAB function fmincon. Optimized negative log likelihoods were compared by means of the sample-size corrected Akaike Information Criterion (AICc, Equation 9 and 10). The model with the lowest AICc value was considered to account best for the observed participants’ choice data, penalized for model complexity and sample size (number of participants).

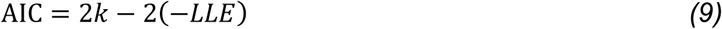

Where k is the number of free parameters of the model and −LLE is the negative log likelihood of the model given the data.

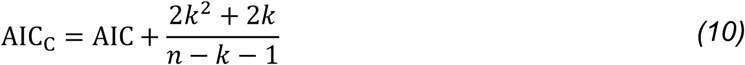

Where k is the number of free parameters in the model, and n is the sample size.

Additionally, we performed 10,000 simulations of choice behavior per participant. After value estimation as described above, using individual parameters of the best-fitting models at group level, the resulting value estimates for each stimulus were entered in the exact same sequence of 120 choices that the participants individually experienced during the decision probe phase. Simulated overall choice probabilities were averaged per participant across 10,000 simulations.

## Acknowledgments

The authors thank all volunteers who participated in this study. The authors would like to thank Halla Mulla-Osman, Stefanie Linnhoff, Nicola Harzen and Denise Scheermann for their invaluable support during data acquisition. We thank the Center for Magnetic Resonance Research of the University of Minnesota for providing the Multiband accelerated fMRI sequence. The authors additionally thank Theo O.J. Gruendler, Lukas M. Neugebauer and Daniel Priegnitz for helpful discussions during set-up of the study and for sharing custom MATLAB code.

## Author contributions

LL designed the study and conceptualized research, acquired the data, analyzed the data, drafted the manuscript, read and edited versions of the manuscript and approved the final version of the manuscript. CT set up the MRI acquisition protocol, read and edited versions of the manuscript and approved the final version of the manuscript. LFK analyzed the data, read and edited versions of the manuscript and approved the final version of the manuscript. GJ designed the study and conceptualized research, analyzed the data, read and edited versions of the manuscript and approved the final version of the manuscript.

## Data and materials availability

The behavioral data, univariate extracted parameters and neural pattern similarity correlation matrices that support the findings of this study are publicly available along with the custom analysis code for behavioral data analyses, univariate fMRI analyses on extracted parameters, multivariate fMRI analyses and brain-behavioral correlational data analyses at GitHub (https://github.com/LLuettgau/revaluation). Neuroimaging data that support the findings of this study will be made available by the corresponding author upon reasonable request.

## Supplementary Materials

**Supplementary Fig. S1.**
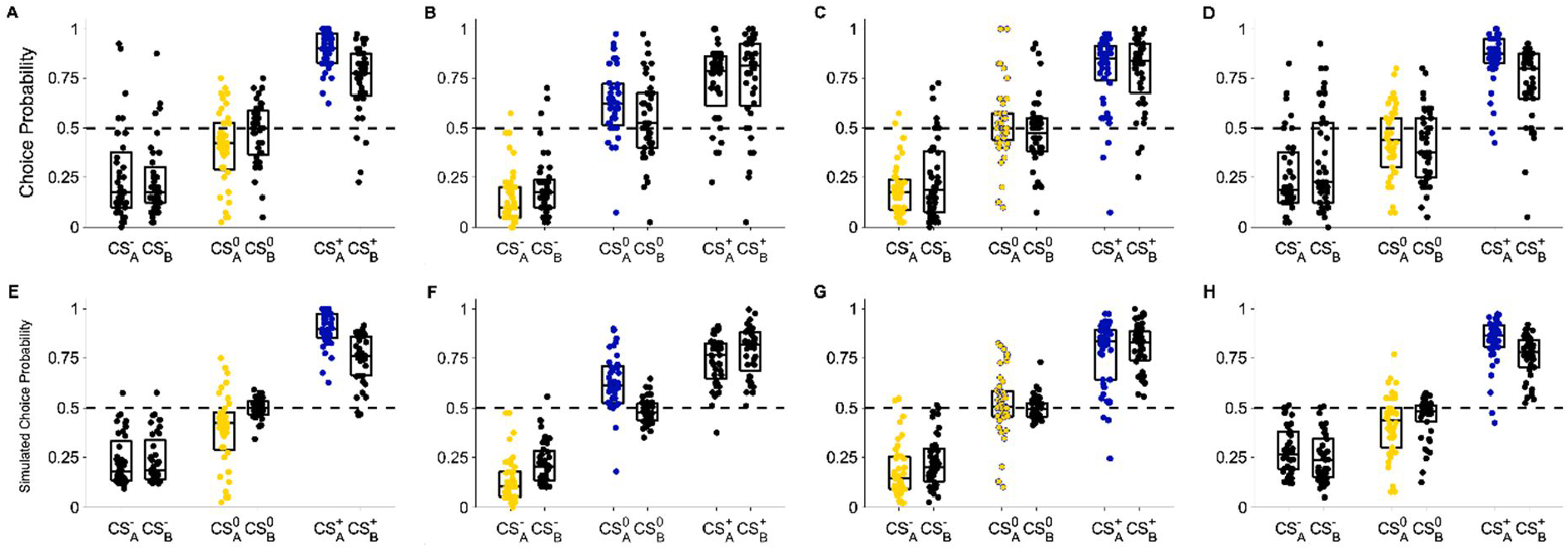
Behavioral and simulation results. A-D) Empirical choice results (as in Fig. 1E-H). Previously chosen CS (blue scatter) are selected more often compared to equivalent CS (black scatter) in Experiments 1, 2, 4 (E, F, H) and previously non-chosen CS (yellow scatter) are selected less often compared to equivalent CS (black scatter) in Experiment 1, 2 (E, F) during decision probe. The effect is not present in Experiment 3 (G), indicating that the roughly equal proportion of choices and non-choices of CS^0^_A_ during revaluation had cancelled each other out. E-H) Averaged simulated choice probabilites (10,000 simulations per participant), recapitulating observed empirical choice patterns.

**Supplementary Fig. S2.**
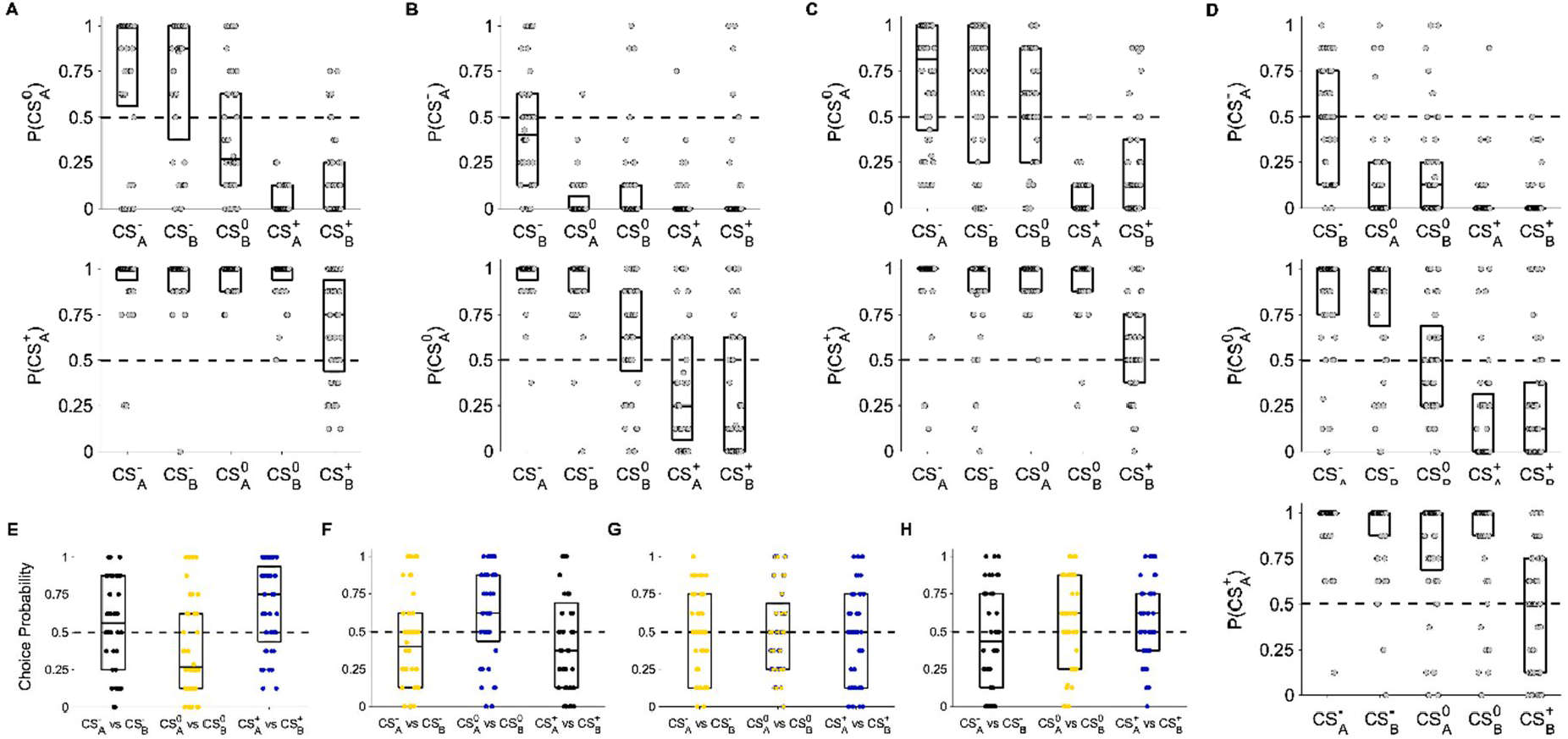
Extended Behavioral Results and Pairwise Choice Probabilities. Decision probe pairwise choice probabilities for CS that were presented during revaluation against every other CS. A) In Experiment 1, CS^0^_A_ is only preferred over CS^-^_A_ and CS^-^_B_ (top), CS^+^_A_ is the most preferred CS (bottom). B) In Experiment 2, CS^-^_A_ is the least preferred CS (top), CS^0^_A_ is chosen more frequently than CS^-^_A_, CS^-^_B_, and CS^0^_B_ (bottom). C) In Experiment 4, CS^0^_A_ is only preferred over CS^-^_A_ and CS^-^_B_ (top), CS^+^_A_ is the most preferred CS (bottom). D) In Experiment 3, CS^0^_A_ is chosen at the same frequency as CS^0^_B_ (middle). E-H) Pairwise within-category choice probabilities displaying the pairwise comparison that are most indicative of choice-induced preference changes. E) Experiment 1, F) Experiment 2, G) Experiment 3, H) Experiment 4. These results suggest that initially conditioned value was not overridden by revaluation choices and that the choice bias was mostly driven by the pairwise decisions of the respective revaluation CS against the same-value CS. However, repetitions of choices between the two revaluation CS also contributed to the observed overall choice probability effects.

**Supplementary Fig. S3.**
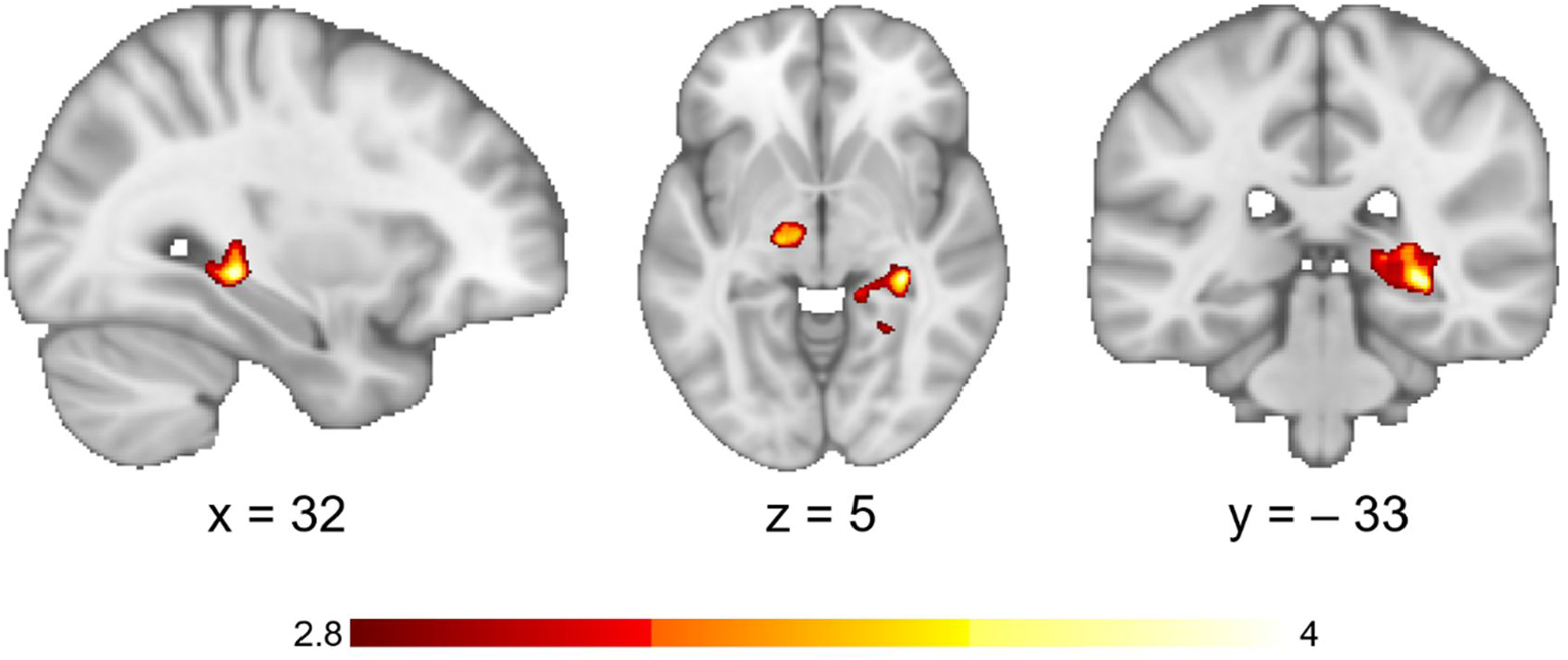
“Cached value” control analysis results. Whole-brain corrected pattern of activation (activation threshold *Z* > 2.3, cluster-forming threshold *P* < 0.05, thresholded at *Z* > 2.8 for display purposes) for the cached value control analysis (Equation 4) during the POST choice-induced revaluation fMRI run. Two clusters in the right posterior hippocampus and in a left midbrain region in the vicinity of the dorsolateral substantia nigra pars compacta survived whole-brain correction. Extracting hippocampal activation with an independent anatomical right hippocampus mask and comparison of parameter estimates for PRE and POST yielded no significant difference (*Z* = 0.91, *P* = .370), suggesting no evidence for an influence of choice-induced revaluation decisions on activation. Additionally, we did not observe significant correlations between the cluster in the right posterior hippocampus from the “cached value” control analysis and the left hippocampus result from the associative analyses (all ρ*s* < .18, *Ps* > .260), suggesting different processes. Parameter estimates from the right posterior hippocampus were only marginally related with overall CP CS^0^_A_ – overall CP CS^0^_B_: ρ_40_ = .30, *P* = .050 (two-tailed) and the within-category CP CS^0^_A_ vs. CS^0^_B_: ρ_40_ = .29, *P* = .062 (two-tailed). However, the direction of the correlation was exactly opposite to the predictions of the cached value account. As there was no apriori anatomical hypothesis for the SNpc, we were unable to extract parameter estimates in an unbiased fashion. Thus, PRE and POST parameter estimates could not be compared. Left SNpc parameter estimates were not significantly related to left VTA parameter estimates from the associative analysis (all ρ*s* < .08, *Ps* > .63), nor with decision probe behavior (all ρ*s* < .20, *Ps* > .20).

**Supplementary Fig. S4.**
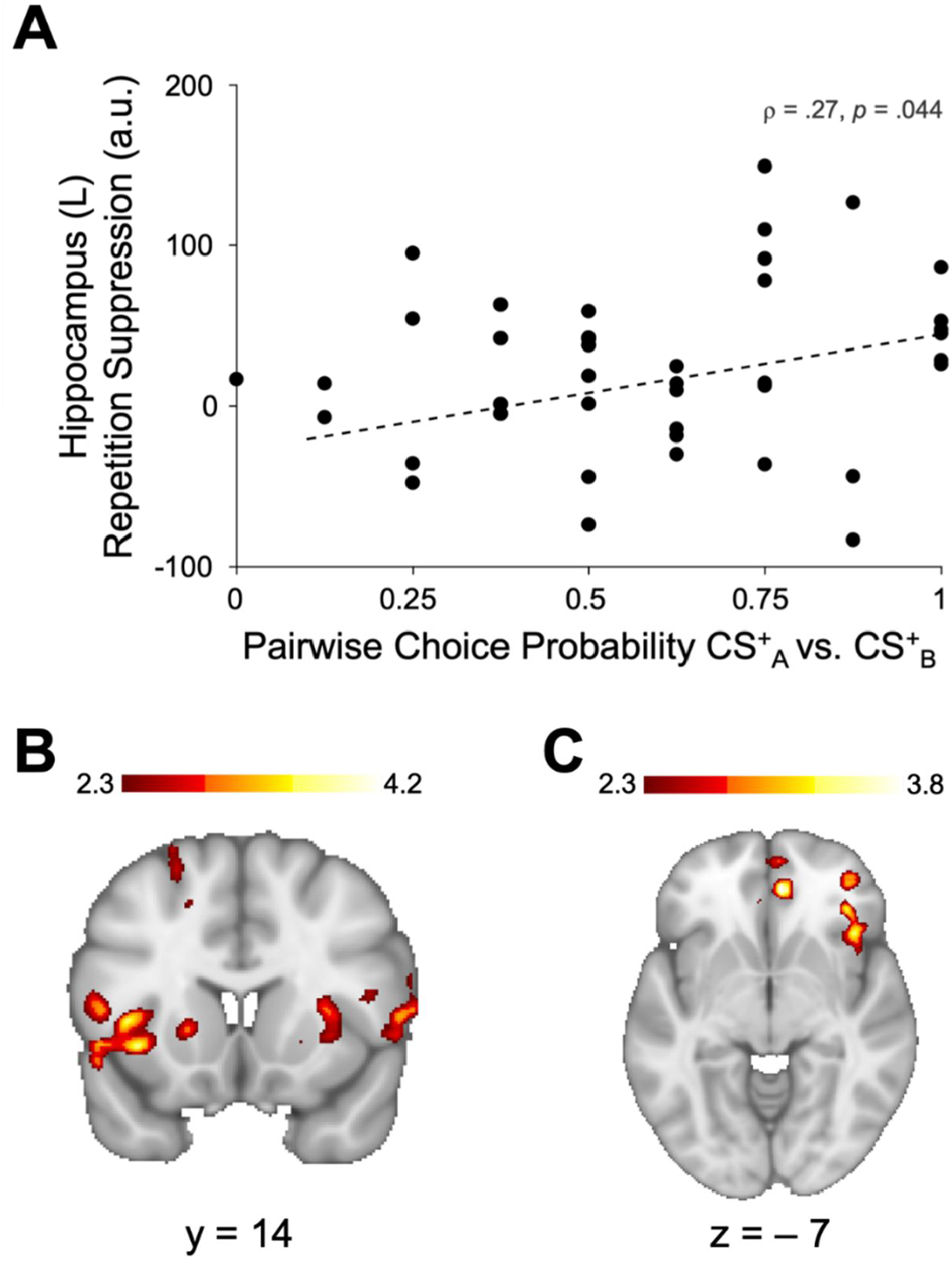
Brain-behavior correlation and whole-brain regressions. A) Left hippocampus parameter estimates (independent anatomical mask, Fig. 4A) for repetition suppression of CS^+^_A_ relative to CS^+^_B_ relative to CS^+^_B_-US^+^, controlling for activation elicited by CS^+^_A_ and CS^+^_B_ followed by both incorrect outcomes (US^-^ and US^0^) (Equation 2) during the POST choice-induced revaluation fMRI run positively correlate with decision probe pairwise choice probability of CS^+^_A_ vs. CS^+^_B_. The higher repetition suppression was after choice-induced revaluation, the more likely participants were to prefer CS^+^_A_ over CS^+^_B_. B) CS^+^_A_ preference vs. CS^+^_B_ was correlated with POST choice-induced revaluation run repetition suppression in the bilateral anterior insula, suggesting choice-induced strengthening of CS-related pre-activation of neural ensembles coding for the used food items. C) We observed a positive relationship between PRE-POST changes in associative strength of CS^0^_A_ and pairwise CP for CS^0^_A_ vs. CS^0^_B_, in two clusters in the medial orbitofrontal cortex and lOFC, extending to the anterior insula.

**Supplementary Table S1.**

| Exp. 1 | Update | Model | $\alpha_{\text{Learning}}$ | $\alpha_{\text{ch}}$ | $\alpha_{\text{unch}}$ | $\tau$ | -LLE | AIC<sub>c</sub> |
| --- | --- | --- | --- | --- | --- | --- | --- | --- |
|  | RW | Model 1 | < .001<br>(0 – 1) | - | - | .0002<br>(0 – 1.55) | 63.43<br>(37.50 – 81.58) | 133.59 |
|  |  | Model 2 | .04<br>(0 – 1) | < .001<br>(0 – .78) | - | .20<br>(0 – 1.48) | 40.52<br>(26.19 – 80.67) | 98.10 |
|  |  | Model 3 | .02<br>(0 – .58) | .50<br>(0 – .85) | .006<br>(0 – .86) | .15<br>(0 – .77) | 37.27<br>(25.62 – 79.88) | **94.62** |
|  | Asso. Value | Model 1 | .002<br>(0 – 1) | - | - | .11<br>(.05 – 2.65) | 41.35<br>(27.38 – 80.9) | 97.10 |
|  |  | Model 2 | .02<br>(0 – 1) | .002<br>(0 – .84) | - | .14<br>(.06 – 1.43) | 40.60<br>(20.64 – 77.37) | 95.87 |
|  |  | Model 3 | .03<br>(0 – 1) | .42<br>(0 – .84) | .005<br>(0 – 1) | .17<br>(.06 – 3.37) | 37.08<br>(20.60 – 81.88) | 96.30 |
| Exp. 2 | RW | Model 1 | .55<br>(0 – .95) | - | - | .45<br>(0 – 66.59) | 53.41<br>(31.86 – 83.18) | 115.99 |
|  |  | Model 2 | .18<br>(0 – .72) | .005<br>(0 – .95) | - | .24<br>(0 – 3.44) | 45.42<br>(15.43 – 82.02) | 103.18 |
|  |  | Model 3 | .04<br>(0 – .85) | .02<br>(0 – .88) | .005<br>(0 – .99) | .24<br>(0 – .77) | 44.67<br>(5.55 – 77.11) | 98.98 |
|  | Asso. Value | Model 1 | .003<br>(0 – 1) | - | - | .15<br>(.05 – 2.65) | 48.77<br>(15.81 – 83.23) | 101.68 |
|  |  | Model 2 | .04<br>(0 – 1) | .005<br>(0 – .92) | - | .20<br>(.002 – 6.11) | 44.53<br>(15.71 – 82.81) | 100.51 |
|  |  | Model 3 | .03<br>(0 – 1) | .02<br>(0 – .99) | .005<br>(0 – .74) | .20<br>(.004 – 2.08) | 44.38<br>(5.55 – 82.4) | **97.43** |
| Exp. 3 | RW | Model 1 | .03<br>(0 – .88) | - | - | .18<br>(0 – 1.55) | 50.60<br>(22.15 – 82.66) | 107.92 |
|  |  | Model 2 | .02<br>(0 – .85) | < .001<br>(0 – .70) | - | .07<br>(0 – 16.92) | 41.06<br>(14.50 – 82.99) | 95.31 |
|  |  | Model 3 | .01<br>(0 – .83) | .009<br>(0 – .55) | < .001<br>(0 – .40) | .08<br>(0 – 24.81) | 40.47<br>(13.64 – 83.07) | 93.15 |
|  | Asso. Value | Model 1 | .005<br>(0 – 1) | - | - | .11<br>(0 – 1.44) | 41.12<br>(13.65 – 82.64) | 93.78 |
|  |  | Model 2 | .02<br>(0 – 1) | .008<br>(0 – .76) | - | .16<br>(0 – 1.28) | 40.29<br>(13.65 – 82.62) | 93.84 |
|  |  | Model 3 | .02<br>(0 – 1) | .012<br>(0 – .86) | .009<br>(0 – .80) | .13<br>(.004 – 2.01) | 44.38<br>(5.55 – 82.4) | **91.96** |
| Exp. 4 | RW | Model 1 | < .001<br>(0 – 1) | - | - | < .001<br>(0 – 3.89) | 71.61<br>(49.60 – 82.72) | 143.94 |
|  |  | Model 2 | < .001<br>(0 – .85) | < .001<br>(0 – 1) | - | .16<br>(0 – 5.05) | 54.25<br>(22.54 – 82.64) | 114.94 |
|  |  | Model 3 | .04<br>(0 – .61) | .52<br>(0 – .85) | .008<br>(0 – .72) | .24<br>(0.03 – 1.02) | 51.11<br>(22.54 – 81.86) | 110.71 |
|  | Asso. Value | Model 1 | .002<br>(0 – 1) | - | - | .16<br>(0.04 – 4.78) | 55.11<br>(23.54 – 82.86) | 112.97 |
|  |  | Model 2 | .13<br>(0 – 1) | .007<br>(0 – 1) | - | .24<br>(0.05 – 4.81) | 50.79<br>(23.37 – 82.61) | 110.69 |
|  |  | Model 3 | .18<br>(0 – 1) | .50<br>(0 – 1) | .009<br>(0 – .97) | .22<br>(.04 – 2.98) | 49.25<br>(22.90 – 82.51) | **107.85** |

*Note:* Median and range of computational parameters, negative log likelihood estimates (–LLE) and corrected Akaike Information Criteria (AIC<sub>c</sub>) of the Rescorla-Wagner like (RW) (Rescorla & Wagner, 1972) and Associative Value (Asso. Value) reinforcement learning models for the four experiments. Lowest AIC<sub>c</sub> per experiment, which was considered to account best for the observed participants' choice data, penalized for model complexity and sample size (number of participants) in bold letters.

